# Explainable Machine Learning Analysis Reveals Gender Differences in the Phenotypic and Neurobiological Markers of Cannabis Use Disorder

**DOI:** 10.1101/2021.08.30.458245

**Authors:** Gregory Niklason, Eric Rawls, Sisi Ma, Erich Kummerfeld, Andrea M. Maxwell, Leyla R. Brucar, Gunner Drossel, Anna Zilverstand

## Abstract

**Background:** Cannabis Use Disorder (CUD) has been linked to environmental, personality, mental health, neurocognitive and neurobiological risk factors. While many studies have revealed gender differences in CUD, the relative importance of these complex factors by gender has not been described.

**Methods:** We conducted a data-driven examination of gender differences in CUD in a community sample of young adults (Human Connectome Project [HCP]; n = 1204, 54% female). We employed state-of-the-art machine learning methods [gradient tree boosting, XGBoost] in combination with novel factor ranking tools [SHapley’s Additive exPlanations (SHAP)] as an ‘explainable machine learning approach’ in the multimodal data collected by the HCP (phenotypic and brain data).

**Results:** We were able to successfully classify both cannabis dependence and cannabis use levels. Previously identified environmental, personality, mental health, neurocognitive, and brain factors highly contributed to the classification. Predominantly-male risk factors included personality (high openness), mental health (high externalizing, high childhood conduct disorder, high fear somaticism), neurocognitive (impulsive delay discounting, slow working memory performance) and brain (low hippocampal volume) factors. Conversely, predominantly-female risk factors included environmental (low education level, low instrumental support) factors.

**Conclusions:** Our data-driven analysis of gender differences in the multimodal risk factors underlying cannabis dependence and use levels demonstrate that environmental factors contribute more strongly to CUD in women, whereas individual factors such as personality, mental health and neurocognitive factors have a larger importance in men. This warrants further investigations, and suggests the importance of understanding how these differences relate to the development of effective treatment approaches.

## 1 Introduction

Cannabis is the most commonly used illicit drug in the United States, with an estimated 8.2% of the population reporting cannabis use in the past month (1). Of those who endorsed past-year use, an estimated 30.6% met criteria for cannabis use disorder (CUD) (2). Previous research has established gender differences in use, however, little is known about the factors that drive these differences (3,4). The factors underlying cannabis use and dependence are complex and have, until now, often only been investigated in a fragmented way, with researchers focusing on a small number of factors in each study. However, the recent availability of large public datasets with broad phenotyping, such as the Human Connectome Project (HCP), and the emergence of novel machine learning approaches and ranking tools for evaluating the importance of each factor (5), make it possible to shift towards an analysis of the broad patterns of factors underlying CUD.

Prominent neurobiological theories of addiction have traditionally focused on the importance of reward- and approach-related behavior, with newer theories integrating cognitive and affective factors as important additional functional domains (6–9). However, even “multi-mechanistic” addiction models are limited, such as the Koob-Volkow model (6) that discusses three main mechanisms involved in addiction: incentive salience/habit formation (reward/approach-related behavior), negative affect, and executive function. Only very few addiction theories have moved beyond this triadic-mechanism framework [e.g. see “vulnerabilities in decision making” (10) as an example], and even less empirical work has been done using a multi-domain data-driven approach [e.g., see “An Integrated Multimodal Model of Alcohol Use Disorder” (11) as an example]. However, separate empirical investigations strongly suggest the involvement of a myriad of different factors.

Individual risk factors that have been shown to predict high likelihood of cannabis abuse and dependence include gender (12), general cognitive ability [IQ/working memory (13–15)], childhood mental health disorders [(depression, externalizing/conduct disorder) (12,16–21)], trauma history (19,22,23) and stressful life events/low socioeconomic status (16). Cannabis users have further been characterized to have personality traits of high openness/extraversion and low agreeableness/conscientiousness, while neuroticism has not been linked to cannabis abuse (24–26). Increased openness in particular appears to discriminate cannabis users from other drug users (26). The triadic neurobiological models of cannabis addiction are supported by evidence on increased reward/approach-related behavior [e.g., increased sensation seeking (27) & delay discounting (28,29)], a role of increased negative affect [e.g. increased prevalence of depression (2,12,16)] and deficits in executive function, specifically deficits in memory/working memory performance and processing speed deficits that predict risk for chronic cannabis use (13–15,30,31). Neuroimaging studies corroborate these theories by demonstrating an upregulation of brain regions involved in reward/approach-related behavior [e.g., salience/reward network (9,32)] and structural changes in valuation networks [e.g., orbitofrontal cortex (33,34)], as well as changes in brain structures supporting memory function [e.g., reduced hippocampal volume (33); altered memory network function (32)]. Finally, reduced educational attainment and lower socioeconomic status have been shown to co-occur with chronic cannabis use (35–37). Specifically, longitudinal studies have concluded that common risk factors [e.g., lack of support in family/peer/school environment (38) and mental health issues (36)] cause both substance use and lower educational attainment/socioeconomic status.

To evaluate the relative importance of a wide variety of factors associated with high cannabis use levels and cannabis dependence as well as potential gender differences in a well described community sample (HCP [39]; N=1204), we employed state-of-the-art machine learning methods [gradient tree boosting, XGBoost (40)] in combination with a novel ranking tool [SHapley’s Additive exPlanations (SHAP) (5)] to assign relative importance (i.e., SHAP values) to each of the associated factors. Decision and boosted tree-based machine learning methods are powerful tools for identifying associated factors in psychiatric research due to their non-parametric nature (resilience to non-normal data distributions) and their tolerance for multicollinear and missing data (41). However, when used on their own, it is difficult to interpret the relative importance of each of the factors involved. We therefore employed SHAP, an extension of methodology originally developed for consistent credit attribution in cooperative game theory (42), to provide a reliable and consistent ranking of the unique relative importance of each factor (5). In addition to providing a ranking for the unique and additive importance of all identified factors, SHAP allows for examining interactions between factors in a model (43). This is particularly relevant for cannabis use, as research in adolescent users indicates that the association of individual risk factors with cannabis use might differ according to gender (4), which has not been studied comprehensively in adults to date (3). In summary, the current study is an exploratory, data-driven analysis that leverages state-of-the-art machine learning algorithms to model the complex factors underlying chronic cannabis use, and their relative importance by gender.

## 2 Methods

### 2.1 Participants

We analyzed data from the final HCP data release [N=1204, aged 22–35, 54% female; https://db.humanconnectome.org/data/projects/HCP_1200; HCP preprocessing pipeline (4.1)]. All participants provided written informed consent at Washington University. In this community sample, 9% of participants met the DSM-IV criteria for cannabis dependence (n = 109, 26% female; note that cannabis abuse was not assessed). See *Table 1* for detailed demographic information.

**Table 1.**
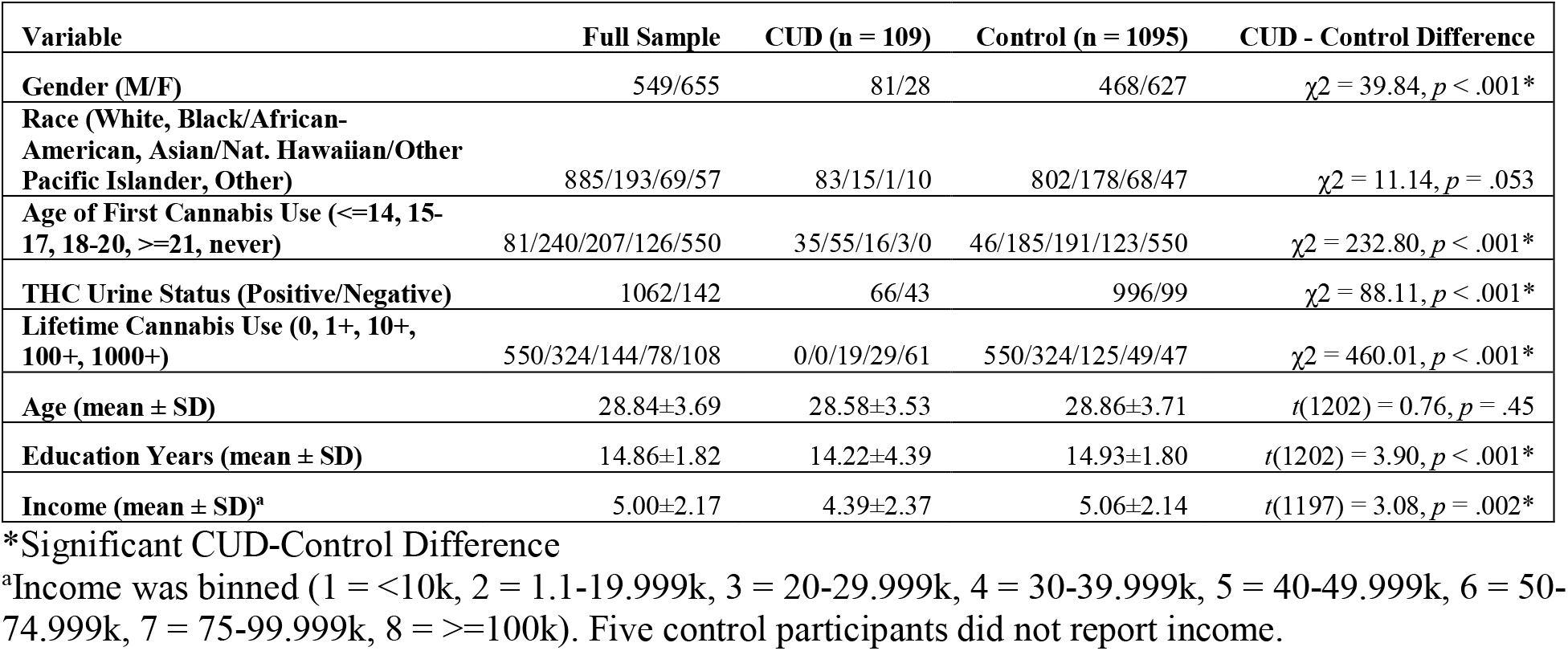
Demographic characteristics of the full sample who completed the SSAGA interview (n = 1204).

### 2.2 Outcome Variables

Our primary outcome measures of interest were 1) level of cannabis use and 2) cannabis dependence, which were assessed using a structured interview (the Semi-Structured Assessment for the Genetics of Alcoholism [SSAGA] (44)). Level of cannabis use was assessed by the reported number of lifetime uses (categories: 0, 1-5, 6-10, 11-100, 101-999, 1000+ lifetime uses). For our analysis, we merged two categories, such that we had five different levels of cannabis use on a logarithmic scale (0, 1+, 10+. 100+, 1000+ lifetime uses). Classification analyses were conducted for each outcome respectively, i.e., we classified escalation of cannabis use and dependence (1+ uses, 10+ uses, 100+ uses, 1000+ uses, and DSM-IV dependence). Each analysis classified a binary outcome using the entire sample; that is, we classified individuals who used cannabis 1+ times from those who did not, classified individuals who used cannabis 10+ times from those who used cannabis <10 times, and so on.

### 2.3 Phenotypic Models

The HCP dataset contains a wide array of self-report, diagnostic and behavioral measures assessing domains of cognition, emotion, social function, psychiatric dysfunction, and personality (45). To examine as broad a phenotypic space as possible, this study used all available behavioral, self-report, and interview-based measures in the HCP database (including all in-scanner task behavior variables). We generally included both summary scores and more specific scores, because the machine learning method we used (detailed below) explicitly allows for correlated factors during model fitting (40). For a complete list of all included phenotypic variables (273 in total), see Supplementary Table 1.

### 2.4 Freesurfer (Structural MRI) Models

For our structural Magnetic Resonance Imaging (MRI) or “Freesurfer” model, we used the Freesurfer data provided by HCP (46,47). This summary data included Freesurfer-generated volume estimates for 44 regions, surface area and cortical thickness estimates for 68 regions, and 19 summary measures including total gray matter volume, white matter volume, and brain segmentation volume (199 factors total).

### 2.5 Resting-State Global and Local Efficiency Models

We used the volumetric resting-state functional MRI (rsfMRI) data as preprocessed by HCP (46). Using the Brain Connectivity Toolbox (48), we conducted a graph theory analysis to extract measures of nodal global and local efficiency [connectivity of a brain region with the rest of the network (global) or with the network within a small neighborhood (local)] from 638 similarly sized brain regions [whole-brain, excluding cerebellum; (49,50) sub-parcellation of the Automated Anatomical Labeling atlas (AAL) (51)]. For each participant, a 638-by-638 matrix of Fisher’s z-transformed Pearson correlations was computed, representing the normalized bivariate correlation of each brain region with each other region. This correlation matrix was binarized at a proportional cost (to improve stability of measures over absolute thresholds) (52) of 0.15 (which is in the middle of the optimal range of 0.01 – 0.30) (53), to represent the strongest 15% of positive connections. We characterized the intrinsic properties of the obtained connectivity graphs by computing nodal global and local efficiency for all 638 brain regions (54), and then averaging both graph theory measures within each larger AAL region (90 factors).

### 2.6 Resting-State Network Connectivity Models

We used the resting-state grayordinate (CIFTI) functional data provided by HCP, to compute within and between functional connectivity for a set of brain networks (55–59). We first parcellated the whole brain into 718 parcels using the Cole-Antecevic parcellation (60–62). We calculated the pairwise Pearson correlations between each pair of parcels in the brain, normalized the obtained correlations using Fisher’s z-transform, and averaged the parcel-to-parcel correlation values both within and between networks (78 factors in total).

### 2.7 Task fMRI Models

All task fMRI (tfMRI) data were preprocessed by HCP using the same steps as for the rsfMRI data (46). We used the provided task fMRI task activation Contrast Of Parameter Estimates (COPE) maps (generated by FSL’s FEAT) that were acquired during seven behavioral tasks, described in (45). These tasks included 1) an N-Back task, 2) a gambling task, 3) a motor mapping task, 4) a language-math task, 5) a social cognition task, 6) a relational-processing task, and 7) an emotion-processing task. We selected 12 COPE maps that represented the main task effects of interest for each task: 1) N-Back task: 2back-0back contrast, 2) gambling task: response to punishments and rewards, 3) motor mapping task: response to left/right foot, left/right hand and tongue movements, 4) language-math task: story-math contrast, 5) social cognition task: social-random contrast, 6) relational-processing task: relational-match contrast, and 7) emotion-processing task: negative faces-shapes contrast. To define activation clusters, we employed the cifti-find-clusters command in Connectome Workbench v1.4.2 (https://www.humanconnectome.org/software/get-connectome-workbench) to find clusters of significantly activated voxels for each of the selected contrast maps, using the full sample (N = 889 with task fMRI). We chose a cutoff of Cohen’s d > 0.8 to select only clusters with large effect sizes and reduce the number of factors entering our final model. Then, for individual participants, we extracted the mean beta weight within each cluster of selected voxels. The task fMRI model contained 448 factors.

### 2.8 Classification Analysis Using Gradient Tree Boosting

To classify each outcome variable of interest, we used a nonparametric classification approach called gradient tree boosting. Gradient tree boosting machines are fit to the gradient of the loss function at every iteration, building up a series of simple models using gradient descent in function space. Specifically, we used the recently developed XGBoost (eXtreme Gradient Boosting) (40), a fast and scalable state-of-the-art gradient tree boosting system. We chose gradient tree boosting because this class of methods is stable and requires a much smaller sample size to produce reliable effect estimates (63), compared to previous methods such as support vector machines (64). For example, XGBoost was able to achieve 95% classification accuracy in a standard benchmarking dataset with a sample size of less than 50 (63).

Nested k-fold cross-validation was used to tune hyperparameters (inner loop) and evaluate classification performance (outer loop) (65). We used k=5, therefore evaluating 5 models using an 80-20 train-test split in both inner and outer loops. We consider the Cartesian product of the following hyperparameters: learning rate={0.01, 0.02, 0.05, 0.1, 0.2}, max tree depth={4, 6, 8, 10, 12}, and subsampling size={0.6, 0.8, 1}. During the inner loop of the nested cross-validation, we conducted a grid search to determine the best combination of the above hyperparameters. The performance of the best model selected from the inner loop was evaluated in the outer loop, resulting in 5 performance estimates. The overall best performing set of hyperparameters for each outcome is reported in the Supplementary Table 2. We additionally used an early stopping parameter of 30 rounds, thus preventing overfitting when the model loss function fails to improve. Since the HCP dataset contains many related participants, our cross-validation scheme always assigned family members to the same group (train or test) for every fold, therefore ensuring that test performance was not inflated by allowing the model to be trained and then tested on a related subject.

We quantified the performance of each model by using the Area Under the Curve of the Receiver Operating Characteristic Curve (AUC-ROCC), which describes how well the model can distinguish between classes. The AUC-ROCC ranges from 0 to 1; higher AUC-ROCCs indicate better predictive performance. An AUC-ROCC of 0.5 indicates random prediction for a binary outcome.

### 2.9 Factor Importance Ranking Using SHapley Additive exPlanations

Advanced machine learning methods such as gradient boosting machines are capable of making highly accurate predictions, but often these predictions come at the expense of interpretability. That is, traditional classification approaches do not allow for an interpretation of the relative importance of the factors involved, as they only evaluate the predictive performance of the entire model. To evaluate the unique relative importance of each model factor (referred to as “features” in machine learning research), we used SHAP (SHapley Additive exPlanations), proposed by (5), as a feature ranking tool. SHAP provides an explanation model that computes the unique and additive importance of each model feature (predictive factor) in determining the final classification result. SHAP is based on the concept of Shapley Values, originally described in (42) as a consistent method to allocate credit to a set of team members for a cooperative outcome. In this case, rather than the consortium consisting of a team of players working toward a common goal, the consortium consists of the set of features (factors) which work toward the common goal of producing the classification output of the model. The impact of each feature on the output of the model is defined as the change in model output when the feature is known, as opposed to unknown. Shapley values are the only currently available feature ranking tool that obeys a specific set of properties (local accuracy, consistency, and missingness [5]), which are considered desirable in explaining the output of a machine learning classification model. An in-depth explanation of the properties of SHAP is beyond the scope of the current paper; for a full explanation of the properties of SHAP, the reasons these properties are desirable, and the equations used to derive the feature importance rankings, please see (5) and (43).

### 2.10 Using SHAP to Investigate Gender Effects in Cannabis Use and Dependence

Critical to our current investigation, SHAP is also able to leverage the assumption of feature additivity to compute interaction effects between sets of two factors in the model (43). SHAP values can provide a rich alternative to traditional partial dependence plots (66). While partial dependence plots only allow for an interpretation of how the output of a model depends on the interaction between two factors, SHAP dependence plots allow for interpreting interaction effects while accounting for both lower- and higher-order interaction effects of all factors in the model. In this study, we leveraged this to investigate gender differences in model factors, as gender was a strong predictor of cannabis outcomes in all models.

## 3. Results

### 3.1 Classification performance

Cross-validated AUC-ROCCs of the six unimodal models we considered returned a wide range of performance indices (*Figure 1a*). The phenotypic model had an average AUC-ROCC of 0.70 over all five outcome measures, and produced the best performance in classifying 1000+ cannabis uses (AUC-ROCC = 0.74). Of the brain models, the best performance was obtained by the Freesurfer (structural MRI) model (average AUC-ROCC = 0.58) and the global efficiency model (average AUC-ROCC = 0.57). The other brain models all performed similarly to each other, and were not considered further (AUC-ROCC range = 0.52-0.53).

**Figure 1.**
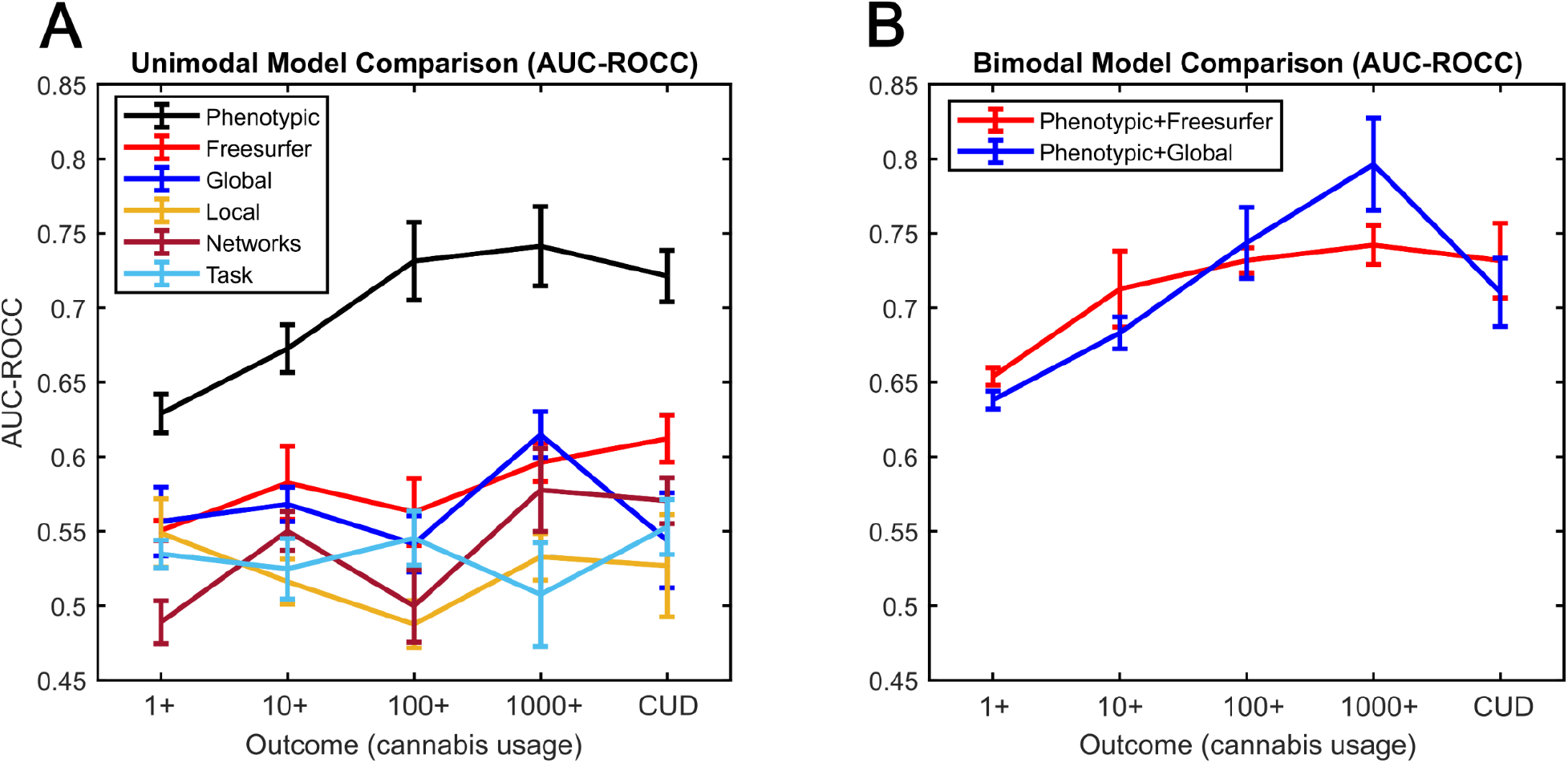
Area under the Curve of the Receiver Operating Characteristic Curve (AUC-ROCC) was used to quantify the classification performance of A: Six unimodal models, and B: Two additional multimodal models that combined the factors of the phenotypic model with the factors of the two best performing brain models. The phenotypic model performed better than any of the other unimodal models, and the inclusion of the two best sets of brain factors (Freesurfer & global efficiency) did not appreciably improve the performance of the phenotypic model. Error bars correspond to ±1 Standard Error.

To determine if performance of the phenotypic model could be improved by adding factors from the most informative brain modalities, we then tested two bimodal models (phenotypic+Freesurfer, phenotypic+global efficiency; *Figure 1b*). For both of the combined models, the average AUC-ROCC over all five outcomes was 0.71. The best performance of the combined models (phenotypic+Freesurfer, phenotypic+global efficiency) was obtained in classifying 1000+ cannabis uses (AUC-ROCC = 0.74 & 0.80, respectively). The results indicate that while the inclusion of brain data did not appreciably change the overall classification accuracy, specific brain factors (e.g. hippocampus volume, median rank = 4) were among the highest ranked predictors in these bimodal models.

### 3.2 SHAP Factor Importance Ranking

To determine which factors drove the performance of the best performing classification models, we used SHAP to estimate the relative importance of all factors (e.g., see the factors contributing to dependence in *Figure 2*; see other models in *Supplementary Figures 1-8*). To determine which factors consistently classified increased cannabis use levels and dependence, we computed the median rank of each factor across all models (see *Table 2*). The consistent highly ranked factors across models (median rank ≤20) included a broad range of factors, such as environmental factors (gender, income, education level), personality measures (openness), mental health measures (externalizing, childhood conduct disorder, aggression), neurocognitive measures (working memory, verbal IQ) and brain measures (hippocampal, brainstem and CSF volume; frontal pole thickness; insula, operculum and occipital resting-state connectivity) (*Table 2*).

**Figure 2.**
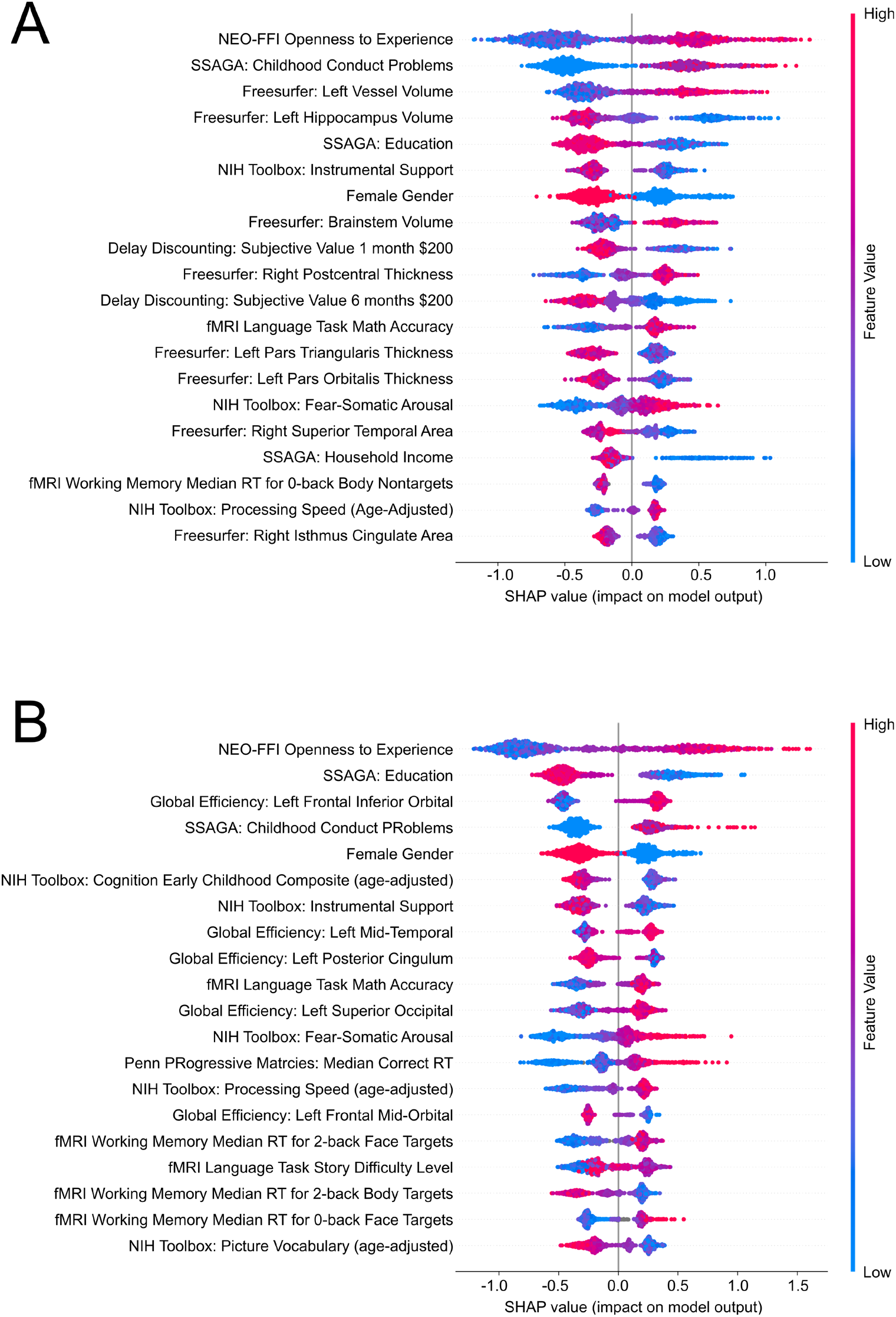
SHAP ranking for factors contributing to the two bimodal models (A: phenotypic+Freesurfer, B: phenotypic+global efficiency) classifying cannabis dependence. Factors are ranked in order of the average (absolute) SHAP value, which indicates the importance of a factor. Individual data points (dots) represent the model output for each individual in the sample. The position of a dot on the x-axis represents the impact of the observed factor on the model output for the individual. More positive SHAP values on the x-axis indicate that the observed factor pushed the classification closer towards cannabis dependence, whereas more negative SHAP values indicate that the factor pushed the classification away from cannabis dependence. The color of the individual dots represent the value of the observed measurements, with blue indicating lower and red higher values (e.g. red indicates high openness or high education level in these plots). As an example, in both plots high (red dots) observed “Openness to Experience” and low (blue dots) education levels pushed the model prediction closer towards cannabis dependence. That is, high “Openness to Experience” and lower education levels make it more likely that the model would categorize a given individual as cannabis-dependent relative to not dependent.

**Table 2.** SHAP factor ranking for the two bimodal classification models. Only factors with a median rank of 20 or less (i.e., the highest ranked among the total included >1000 features) are presented here.

| Factor | Rank<br>1+ | Rank<br>10+ | Rank<br>100+ | Rank<br>1000+ | Rank<br>CUD | Median<br>Rank |
| --- | --- | --- | --- | --- | --- | --- |
| *Phenotypic + Freesurfer Model* |  |  |  |  |  |  |
| NEO-FFI Openness | 1 | 1 | 1 | 1 | 1 | 1 |
| ASR Externalizing | 2 | 6 | 2 | 2 | 70 | 2 |
| Freesurfer: Left Hippocampus Volume | 4 | 3 | 8 | 52 | 4 | 4 |
| Female Gender | 89 | 2 | 3 | 4 | 7 | 4 |
| NIH Toolbox Anger Aggression | 7 | 4 | 6 | 8 | 75 | 7 |
| Freesurfer: Brainstem Volume | 6 | 5 | 110 | 48 | 8 | 8 |
| Freesurfer: CSF Volume | 8 | 11 | 10 | 10 | 110 | 10 |
| Freesurfer: Left Frontal Pole Thickness | 44 | 9 | 14 | 6 | 87 | 14 |
| fMRI N-Back Task RT: 2-back place target | 12 | 119 | 15 | 14 | 53 | 15 |
| fMRI N-Back Task RT: 2-back body target | 10 | 16 | 20 | 16 | 22 | 16 |
| SSAGA Income | 270 | 12 | 5 | 19 | 17 | 17 |
| SSAGA Childhood Conduct | 18 | 21 | 17 | 12 | 2 | 17 |
| NIH Toolbox Picture Vocabulary (age-adjusted) | 13 | 32 | 19 | 13 | 41 | 19 |
| *Phenotypic + Global Efficiency Model* |  |  |  |  |  |  |
| NEO-FFI Openness | 1 | 1 | 1 | 2 | 1 | 1 |
| ASR Externalizing | 2 | 3 | 4 | 6 | 35 | 4 |
| Female Gender | 121 | 4 | 3 | 3 | 5 | 4 |
| fMRI N-Back Task RT: 2-back body target | 5 | 8 | 6 | 12 | 18 | 8 |
| NIH Toolbox Picture Vocabulary (age-adjusted) | 8 | 6 | 7 | 42 | 20 | 8 |
| NIH Toolbox Anger Aggression | 6 | 2 | 13 | 13 | 27 | 13 |
| SSAGA Education | 246 | 14 | 2 | 33 | 2 | 14 |
| SSAGA Income | 165 | 11 | 8 | 15 | 52 | 15 |
| Global Efficiency: Left Rolandic Operculum | 10 | 13 | 15 | 32 | 90 | 15 |
| fMRI N-Back Task RT: 2-back body nontarget | 102 | 16 | 10 | 18 | 85 | 18 |
| Global Efficiency: Left Insula | 9 | 18 | 105 | 19 | 151 | 19 |
| Global Efficiency: Left Superior Occipital | 19 | 12 | 32 | 25 | 11 | 19 |
| fMRI N-Back Task RT: 2-back place target | 11 | 75 | 12 | 20 | 40 | 20 |

### 3.3 SHAP Gender Interaction Analysis

Since gender was a top ranked factor (ranked 4th across all phenotypic+Freesurfer and phenotypic+Global models, Table 2), we examined interaction effects to identify gender-specific factors that contribute to classifying cannabis dependence. We focused on the models predicting cannabis dependence and use levels of 1000+ lifetime uses as the most clinically relevant outcomes. We report all interaction effects with a SHAP interaction value of at least 0.1 (the sum of all SHAP values per model is 1), in order to discuss only interaction effects with meaningful effect sizes.

#### 3.3.1 SHAP Gender Interactions in Models Predicting Cannabis Dependence

The bimodal models (phenotypic+Freesurfer; phenotypic+global) classifying cannabis dependence indicated gender interaction effects for environmental factors (education level), personality measures (openness), mental health factors (childhood conduct disorder, fear somaticism), neurocognitive measures (delay discounting, working memory) and brain measures (hippocampal volume, postcentral thickness, superior temporal area) (*Figures 3+4*). Males as compared to females were more often classified as cannabis-dependent based on personality (high openness), mental health (high childhood conduct disorder, high fear somaticism), neurocognitive (impulsive delay discounting, slow working memory performance) and brain factors (low hippocampal volume, high postcentral thickness). In contrast, females were more often classified as dependent based on environmental (lower education level) and brain factors (smaller superior temporal area).

**Figure 3.**
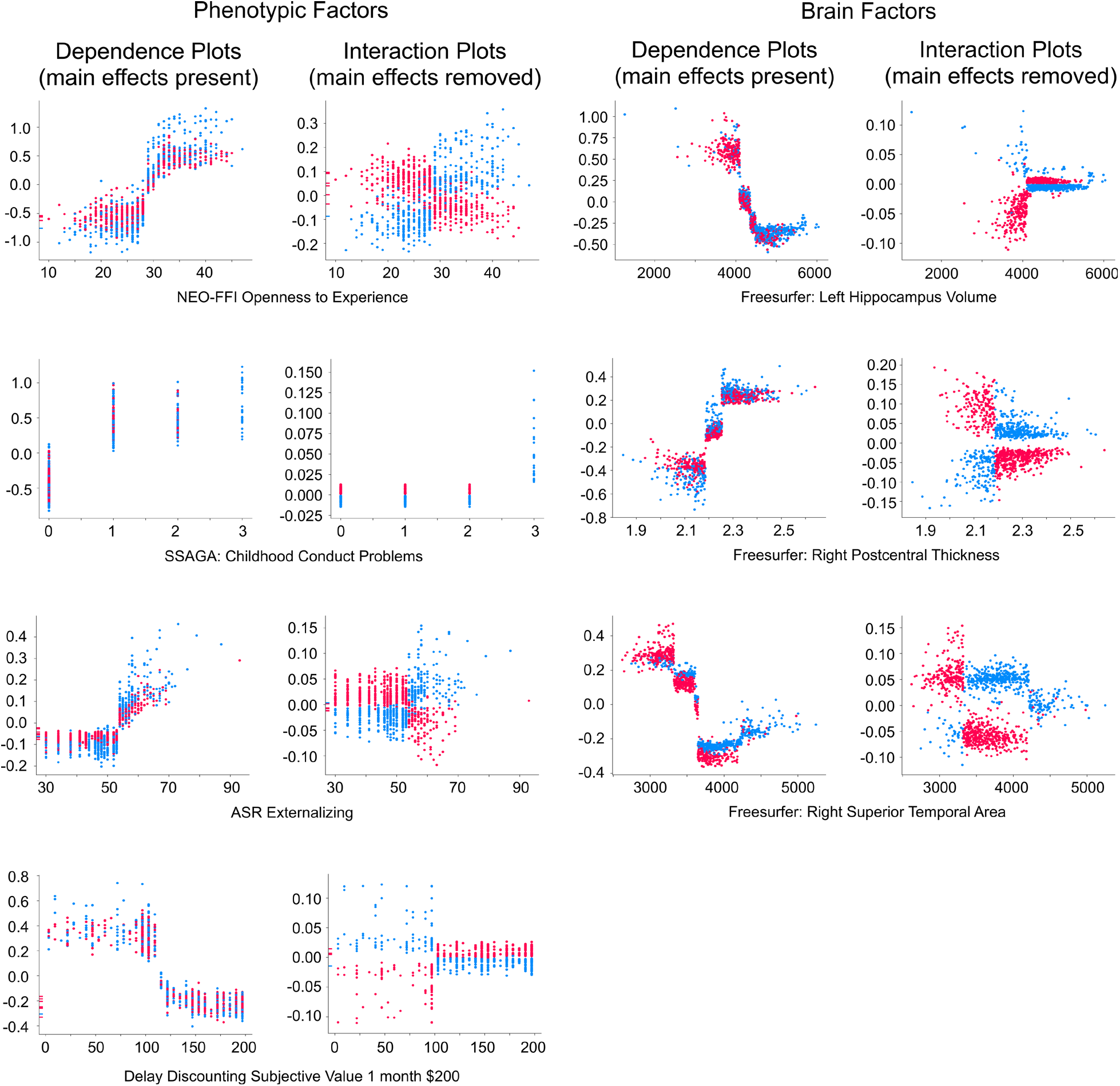
Several factors in the phenotypic+Freesurfer model classifying cannabis dependence showed gender interaction effects. As in previous figures, each dot indicates an individual. Dots are colored according to gender (red = female, blue = male). The left columns (“Dependence plots”) show individuals colored according to gender, with main effects intact; the right columns (“Interaction plots”) again show the same individuals, but with main effects removed for improved visualization of the gender interaction effects. The x-axis of the plots indicates the observed measurement value of each factor, and the y-axis of each plot indicates the SHAP value (where a higher SHAP value pushed the model closer towards dependence, and lower values pushed the model away from dependence).

**Figure 4.**
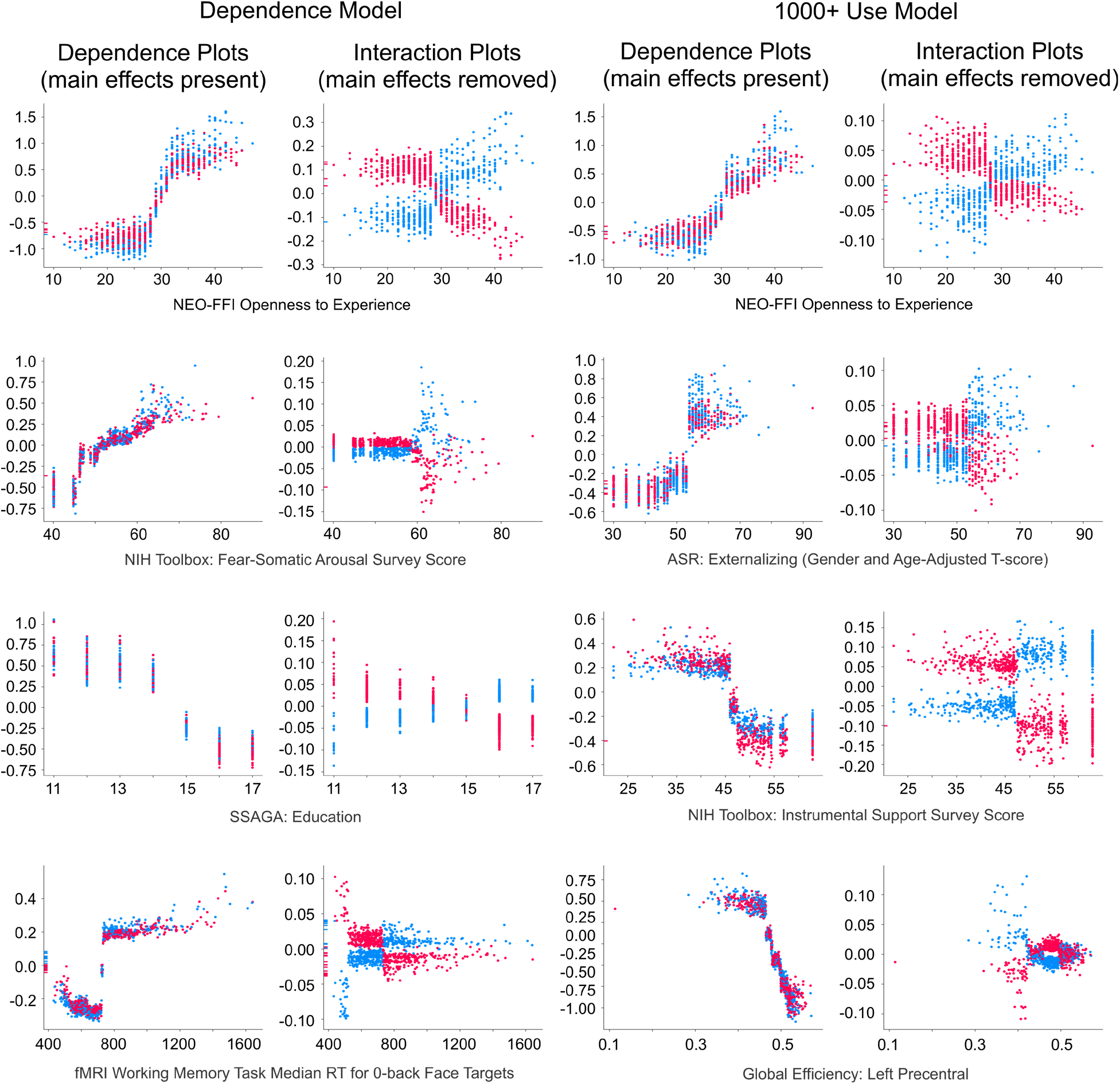
Several factors in the phenotypic+Global Efficiency model classifying cannabis dependence (left) and classifying heavy cannabis use (1000+ uses; right) showed gender interaction effects. As in previous figures, each dot indicates an individual. Dots are colored according to gender (red = female, blue = male). The left columns (“Dependence plots”) show individuals colored according to gender, with main effects intact; the right columns (“Interaction plots”) again show the same individuals, but with main effects removed for improved visualization of gender interaction effects. The x-axis of the plots indicates the observed measurement value of each factor, and the y-axis of each plot indicates the SHAP value (where a higher SHAP value pushed the model closer towards dependence, and lower values pushed the model away from dependence).

#### 3.3.2 SHAP Gender Interactions in Models Predicting Heavy Cannabis Use

The 1000+ lifetime uses model demonstrated gender interaction effects for environmental factors (instrumental support), personality measures (openness), mental health factors (externalizing) and brain measures (precentral efficiency) (see *Figure 4* for gender interactions in the phenotypic+global model; the phenotyopic+Freesurfer model showed no gender interaction effects >0.1). Males as compared to females were more often classified as heavy cannabis users (+1000 uses) based on personality (high openness), mental health (high externalizing) and brain factors (low global efficiency of the precentral cortex). In contrast, females were more often classified as heavy cannabis users based on environmental factors (low instrumental support).

## 4 Discussion

The current study used a machine learning approach in a community sample of young adults to describe the complex factors underlying chronic cannabis use and their relative importance by gender. While a number of recent reviews have recognized the potential for machine learning methods in psychiatric research (67–72), this is the first study to date to use such an approach in adults with CUD, although machine learning methods have been applied to examine adolescent cannabis use (4). Therefore, it is also the first study to date to comprehensively study gender differences in CUD in adults. Since conventional machine learning methods obtain increased predictive power at the cost of interpretability (69,73,74), we paired our classification models [(XGBoost) (40)] with SHapley Additive eXplanations [(SHAP) (5)] to generate “explainable” machine learning models that rank factors (synonymously called “features” in machine learning research) according to their unique and additive importance in classifying an outcome.

Overall, the classification models achieved high accuracy, which in itself was remarkable, since the used dataset was not designed to assess substance use and dependence (see Rawls and colleagues [11] for a more in-depth discussion of the assessments and how they relate to addiction). The current results further confirmed that a small number of factors of the more than one thousand included model factors, consistently provided a unique and additive contribution to the classification performance, beyond other factors in the model. The identified factors included environmental, personality, mental health, neurocognitive and brain measures, demonstrating the complexity of the factors involved in CUD. Overall, the current results confirm the importance of multi-domain investigations into the factors underlying drug addiction, as in our previous empirical investigation of multi-domain factors in substance use disorders (11).

Many factors that have been well described in the literature on CUD were replicated in this study. The environmental factors that most consistently contributed highly to model classification performance were gender, income and education level. Previous studies have often linked male gender to a higher prevalence of CUD (3,4,12). Previous longitudinal research further suggests that reduced educational attainment and lower socioeconomic status co-occur with (but do not directly cause) chronic cannabis abuse and dependence (35–38). The current results further replicate previous work that has linked the personality trait openness to high cannabis use levels and dependence, suggesting that high openness is a predictor specifically for cannabis as a primary drug of choice (24–26). The current results also confirm an important role of externalizing mental health disorders, aggression and a history of child conduct disorder, which have all been identified as risk factors for cannabis abuse and dependence in longitudinal research (17–21). Notably, while our results provide additional support for an important role of externalizing disorders (e.g. [19–21]), we could not confirm a link between cannabis abuse or dependence and internalizing disorders, as had been reported by some other studies (e.g. [12,16]). Further, in the current study, working memory and verbal IQ measures were among the most highly ranked neurocognitive factors, both of which have consistently been associated with CUD and shown to be risk factors for (not consequences of) cannabis abuse and dependence (13–15,30,31). Finally, brain measures that were consistently highly ranked included hippocampal volume, an important structure of the brain’s memory system (32,75,76), as well as brainstem volume, frontal pole thickness, insula, operculum and occipital resting-state connectivity, all of which are part of the reward, salience and visual brain networks that are most densely innervated by dopaminergic receptors (77). These results converge with previous studies and systematic reviews that have demonstrated that CUD is characterized by changes in the brain’s memory system (32,33,78), the reward and salience networks (32,34), and the occipital lobe (79,80). These results also demonstrate changes in the brain’s reward/approach-related system, a domain that was not captured well by the behavioral assessments or neuroimaging tasks used in this study (see Rawls and colleagues [11] for a more in depth discussion). Thus, the current evidence supports the triadic models of cannabis addiction by indicating changes in the brain’s reward/approach system, deficits in executive function, specifically in working memory function and verbal IQ, and a role of negative affect, specifically of externalizing symptoms and aggression.

The analysis of gender interaction effects revealed complex gender differences in the multi-domain factors underlying cannabis abuse and dependence. Environmental factors such as educational attainment and instrumental support (the latter was not among the highest ranked factors overall) were factors that primarily contributed to model prediction accuracy in female individuals. In stark contrast to this finding, ‘classic’ personality, mental health and neurocognitive factors that have often been linked to chronic cannabis use and dependence in previous studies were primarily driving effects in male individuals. In particular, the ‘male-dominated’ factors included the personality trait openness, a history of conduct disorder, externalizing symptoms, and working memory performance. For brain factors, there were both ‘female-dominated’ factors, such as structural changes in the temporal lobe (which was not among the highest ranked overall factors), and ‘male-dominated’ factors, such as low hippocampal volume and changes in the somatosensory-motor system. In short, these results suggest that brain factors contribute to cannabis use levels and dependence in either gender, whereas environmental factors (educational attainment, instrumental support) play a larger role in females and the ‘classic’ individual factors that have been most often linked to cannabis addiction, contribute more strongly to CUD in males.

The current results provide compelling evidence for gender differences in the multifactorial factors underlying CUD in adults, which had not been previously investigated using a multi-domain approach. We are only aware of one previous study on gender differences in CUD in adults (81). This study specifically investigated gender differences in the role of social support and found a stronger protective relationship of social support in women as compared to men (81). Additionally, our results extend previous findings on cannabis use in adolescence that suggest a stronger influence of environmental factors in girls as compared to boys (82–85). A twin study found that the overall contribution of environmental factors for predicting cannabis use levels, as compared to individual predictive factors, was larger in adolescent girls versus boys (82). Similarly, a longitudinal study described that environmental influences such as attending public (versus private) schools, academic performance, living in a single-parent family, spending time in bars/discos and drug use among friends had a stronger influence on cannabis use levels in adolescent girls as compared to boys (83). The same study found that individual factors such as prior history of smoking/alcohol consumption and antisocial behavior were stronger predictors in adolescent boys (83). Furthermore, one study demonstrated that a protective family environment had a stronger influence on cannabis use onset in adolescent girls as compared to boys (84), and that higher life satisfaction was a stronger protective factor against frequent cannabis use among adolescent girls than boys (85). Overall, the resemblance of the general pattern of a stronger influence of environmental versus individual factors in females in adolescence and adulthood is striking and warrants further investigation.

## Conclusion

In summary, our data-driven investigation of the underlying factors of CUD in adults revealed that a small number of environmental, personality, mental health, neurocognitive and brain factors were consistently linked to cannabis use levels and dependence. These findings largely replicated previous research, and additionally demonstrate that environmental factors contribute more strongly to CUD in women, whereas individual factors such as personality, mental health and neurocognitive factors have a larger importance in men. The current findings therefore warrant further investigations into gender differences in adults with CUD and suggest the importance of understanding how these differences relate to the development of effective treatment approaches.

## Acknowledgments

ER is supported by a postdoctoral training grant from the National Institutes of Mental Health (NIMH; T32 MH115866). GD is supported by a predoctoral training grant from the National Institute on Drug Addiction (NIDA; T32 DA007234). AMM is supported by a predoctoral training grant from the National Institute of Neurological Disorders and Stroke (NINDS; T32 NS105604-04). EK received support for this work from the National Center for Advancing Translational Sciences of the National Institutes of Health Award Number (NCATS; UL1TR000114). The content of this manuscript is solely the responsibility of the authors and does not necessarily reflect the views of NIMH, NINDS, NIDA, or NCATS.

## References

1. SAMSHA. Key Substance Use and Mental Health Indicators in the United States: Results from the 2018 National Survey on Drug Use and Health. 2018;82.

2. Hasin DS, Saha TD, Kerridge BT, Goldstein RB, Chou SP, Zhang H, et al. Prevalence of Marijuana Use Disorders in the United States Between 2001-2002 and 2012-2013. JAMA Psychiatry. 2015 Dec 1;72(12):1235–42.

3. Greaves L, Hemsing N. Sex and Gender Interactions on the Use and Impact of Recreational Cannabis. Int J Environ Res Public Health. 2020 Jan 14;17(2):E509.

4. Spechler PA, Allgaier N, Chaarani B, Whelan R, Watts R, Orr C, et al. The initiation of cannabis use in adolescence is predicted by sex-specific psychosocial and neurobiological features. Eur J Neurosci. 2019;50(3):2346–56.

5. Lundberg SM, Lee S-I. A Unified Approach to Interpreting Model Predictions. In: Advances in Neural Information Processing Systems. Curran Associates, Inc.; 2017. p. 4765–74.

6. Koob GF, Volkow ND. Neurobiology of addiction: a neurocircuitry analysis. Lancet Psychiatry. 2016 Aug;3(8):760–73.

7. Bickel WK, Mellis AM, Snider SE, Athamneh LN, Stein JS, Pope DA. 21st century neurobehavioral theories of decision making in addiction: Review and evaluation. Pharmacol Biochem Behav. 2018 Jan;164:4–21.

8. Yücel M, Oldenhof E, Ahmed SH, Belin D, Billieux J, Bowden-Jones H, et al. A transdiagnostic dimensional approach towards a neuropsychological assessment for addiction: an international Delphi consensus study. Addiction. 2019;114(6):1095–109.

9. Zilverstand A, Goldstein RZ. Chapter 3 - Dual models of drug addiction: the impaired response inhibition and salience attribution model. In: Verdejo-Garcia A, editor. Cognition and Addiction [Internet]. Academic Press; 2020 [cited 2021 Jan 13]. p. 17–23. Available from: http://www.sciencedirect.com/science/article/pii/B9780128152980000034

10. Redish AD, Jensen S, Johnson A. A unified framework for addiction: Vulnerabilities in the decision process. Behav Brain Sci. 2008 Aug;31(4):415–87.

11. Rawls E, Kummerfeld E, Zilverstand A. An integrated multimodal model of alcohol use disorder generated by data-driven causal discovery analysis. Commun Biol. 2021 Mar 31;4(1): 1–12.

12. Meier MH, Hall W, Caspi A, Belsky DW, Cerdá M, Harrington HL, et al. Which adolescents develop persistent substance dependence in adulthood? Using population-representative longitudinal data to inform universal risk assessment. Psychol Med. 2016 Mar;46(4):877–89.

13. Khurana A, Romer D, Betancourt LM, Hurt H. Working Memory Ability and Early Drug Use Progression as Predictors of Adolescent Substance Use Disorders. Addict Abingdon Engl. 2017 Jul;112(7):1220–8.

14. Wilson S, Malone SM, Venables NC, McGue M, Iacono WG. Multimodal indicators of risk for and consequences of substance use disorders: Executive functions and trait disconstraint assessed from preadolescence into early adulthood. Int J Psychophysiol Off J Int Organ Psychophysiol. 2019 Dec 19;

15. Meier MH, Caspi A, Danese A, Fisher HL, Houts R, Arseneault L, et al. Associations between adolescent cannabis use and neuropsychological decline: a longitudinal co-twin control study. Addict Abingdon Engl. 2018 Feb;113(2):257–65.

16. Schlossarek S, Kempkensteffen J, Reimer J, Verthein U. Psychosocial Determinants of Cannabis Dependence: A Systematic Review of the Literature. Eur Addict Res. 2016;22(3):131–44.

17. Defoe IN, Khurana A, Betancourt L, Hurt H, Romer D. Disentangling longitudinal relations between youth cannabis use, peer cannabis use, and conduct problems: developmental cascading links to cannabis use disorder. Addiction. 2019;

18. Pingault J-B, Côté SM, Galéra C, Genolini C, Falissard B, Vitaro F, et al. Childhood trajectories of inattention, hyperactivity and oppositional behaviors and prediction of substance abuse/dependence: a 15-year longitudinal population-based study. Mol Psychiatry. 2013 Jul;18(7):806–12.

19. Oshri A, Rogosch FA, Burnette ML, Cicchetti D. Developmental pathways to adolescent cannabis abuse and dependence: Child maltreatment, emerging personality, and internalizing versus externalizing psychopathology. Psychol Addict Behav. 2011;25(4):634–44.

20. Griffith-Lendering MFH, Huijbregts SCJ, Mooijaart A, Vollebergh WAM, Swaab H. Cannabis use and development of externalizing and internalizing behaviour problems in early adolescence: A TRAILS study. Drug Alcohol Depend. 2011 Jul 1;116(1):11–7.

21. Farmer RF, Seeley JR, Kosty DB, Gau JM, Duncan SC, Lynskey MT, et al. Internalizing and externalizing psychopathology as predictors of cannabis use disorder onset during adolescence and early adulthood. Psychol Addict Behav. 2015;29(3):541.

22. Mills R, Kisely S, Alati R, Strathearn L, Najman JM. Child maltreatment and cannabis use in young adulthood: a birth cohort study. Addiction. 2017;112(3):494–501.

23. Proctor LJ, Lewis T, Roesch S, Thompson R, Litrownik AJ, English D, et al. Child maltreatment and age of alcohol and marijuana initiation in high-risk youth. Addict Behav. 2017 Dec;75:64–9.

24. Fridberg DJ, Vollmer JM, O’Donnell BF, Skosnik PD. Cannabis users differ from non-users on measures of personality and schizotypy. Psychiatry Res. 2011 Mar 30;186(1):46–52.

25. Ketcherside A, Jeon-Slaughter H, Baine JL, Filbey FM. Discriminability of Personality Profiles in Isolated and Co-Morbid Marijuana and Nicotine Users. Psychiatry Res. 2016 Apr 30;238:356–62.

26. Terracciano A, Löckenhoff CE, Crum RM, Bienvenu OJ, Costa PT. Five-Factor Model personality profiles of drug users. BMC Psychiatry. 2008 Apr 11;8(1):22.

27. Creemers HE, van Lier PAC, Vollebergh WAM, Ormel J, Verhulst FC, Huizink AC. Predicting onset of cannabis use in early adolescence: the interrelation between high-intensity pleasure and disruptive behavior. The TRAILS Study. J Stud Alcohol Drugs. 2009 Nov;70(6):850–8.

28. Amlung M, Vedelago L, Acker J, Balodis I, MacKillop J. Steep Delay Discounting and Addictive Behavior: A Meta-Analysis of Continuous Associations. Addict Abingdon Engl. 2017 Jan;112(1):51–62.

29. Strickland JC, Lee DC, Vandrey R, Johnson MW. A systematic review and meta-analysis of delay discounting and cannabis use. Exp Clin Psychopharmacol. 2020 Apr 20;

30. Meier MH, Caspi A, Ambler A, Harrington H, Houts R, Keefe RSE, et al. Persistent cannabis users show neuropsychological decline from childhood to midlife. Proc Natl Acad Sci. 2012 Oct 2;109(40):E2657–64.

31. Gonzalez R, Pacheco-Colón I, Duperrouzel JC, Hawes SW. Does Cannabis Use Cause Declines in Neuropsychological Functioning? A Review of Longitudinal Studies. J Int Neuropsychol Soc JINS. 2017 Oct;23(9–10):893–902.

32. Zilverstand A, Huang AS, Alia-Klein N, Goldstein RZ. Neuroimaging Impaired Response Inhibition and Salience Attribution in Human Drug Addiction: A Systematic Review. Neuron. 2018 Jun;98(5):886–903.

33. Lorenzetti V, Chye Y, Silva P, Solowij N, Roberts CA. Does regular cannabis use affect neuroanatomy? An updated systematic review and meta-analysis of structural neuroimaging studies. Eur Arch Psychiatry Clin Neurosci. 2019 Feb 1;269(1):59–71.

34. Batalla A, Bhattacharyya S, Yücel M, Fusar-Poli P, Crippa JA, Nogué S, et al. Structural and Functional Imaging Studies in Chronic Cannabis Users: A Systematic Review of Adolescent and Adult Findings. PLOS ONE. 2013 Feb 4;8(2):e55821.

35. Maggs JL, Staff J, Kloska DD, Patrick ME, O’Malley PM, Schulenberg J. Predicting Young Adult Degree Attainment by Late Adolescent Marijuana Use. J Adolesc Health Off Publ Soc Adolesc Med. 2015 Aug;57(2):205–11.

36. Danielsson A-K, Falkstedt D, Hemmingsson T, Allebeck P, Agardh E. Cannabis use among Swedish men in adolescence and the risk of adverse life course outcomes: results from a 20 year-follow-up study. Addict Abingdon Engl. 2015 Nov;110(11): 1794–802.

37. Green KM, Doherty EE, Ensminger ME. Long-term consequences of adolescent cannabis use: Examining intermediary processes. Am J Drug Alcohol Abuse. 2017 Sep;43(5):567–75.

38. Verweij KJH, Huizink AC, Agrawal A, Martin NG, Lynskey MT. Is the relationship between early-onset cannabis use and educational attainment causal or due to common liability? Drug Alcohol Depend. 2013 Dec 1;133(2):580–6.

39. Van Essen DC, Smith SM, Barch DM, Behrens TEJ, Yacoub E, Ugurbil K. The WU-Minn Human Connectome Project: An overview. NeuroImage. 2013 Oct;80:62–79.

40. Chen T, Guestrin C. XGBoost: A Scalable Tree Boosting System. In: Proceedings of the 22nd ACM SIGKDD International Conference on Knowledge Discovery and Data Mining [Internet]. New York, NY, USA: Association for Computing Machinery; 2016 [cited 2020 Sep 8]. p. 785–94. (KDD’16). Available from: https://doi.org/10.1145/2939672.2939785

41. Song Y, Lu Y. Decision tree methods: applications for classification and prediction. Shanghai Arch Psychiatry. 2015 Apr 25;27(2):130–5.

42. Shapley LS. A Value for N-Person Games. In: Contributions to the Theory of Games. 2nd ed. Princeton University Press; 1953. p. 307–17.

43. Lundberg SM, Erion GG, Lee S-I. Consistent Individualized Feature Attribution for Tree Ensembles. In: arXiv. 2019.

44. Hesselbrock M, Easton C, Bucholz KK, Schuckit M, Hesselbrock V. A validity study of the SSAGA-a comparison with the SCAN. Addiction. 1999;94(9):1361–70.

45. Barch DM, Burgess GC, Harms MP, Petersen SE, Schlaggar BL, Corbetta M, et al. Function in the human connectome: Task-fMRI and individual differences in behavior. NeuroImage. 2013 Oct;80:169–89.

46. Glasser MF, Sotiropoulos SN, Wilson JA, Coalson TS, Fischl B, Andersson JL, et al. The minimal preprocessing pipelines for the Human Connectome Project. NeuroImage. 2013 Oct;80:105–24.

47. Uğurbil K, Xu J, Auerbach EJ, Moeller S, Vu AT, Duarte-Carvajalino JM, et al. Pushing spatial and temporal resolution for functional and diffusion MRI in the Human Connectome Project. NeuroImage. 2013 Oct;80:80–104.

48. Rubinov M, Sporns O. Complex network measures of brain connectivity: Uses and interpretations. NeuroImage. 2010 Sep 1;52(3):1059–69.

49. Crossley NA, Mechelli A, Vértes PE, Winton-Brown TT, Patel AX, Ginestet CE, et al. Cognitive relevance of the community structure of the human brain functional coactivation network. Proc Natl Acad Sci U S A. 2013 Jul 9;110(28):11583–8.

50. Zalesky A, Fornito A, Bullmore ET. Network-based statistic: identifying differences in brain networks. NeuroImage. 2010 Dec;53(4):1197–207.

51. Tzourio-Mazoyer N, Landeau B, Papathanassiou D, Crivello F, Etard O, Delcroix N, et al. Automated Anatomical Labeling of Activations in SPM Using a Macroscopic Anatomical Parcellation of the MNI MRI Single-Subject Brain. NeuroImage. 2002 Jan;15(1):273–89.

52. Garrison KA, Scheinost D, Finn ES, Shen X, Constable RT. The (in)stability of functional brain network measures across thresholds. NeuroImage. 2015 Sep;118:651–61.

53. Bullmore ET, Bassett DS. Brain graphs: graphical models of the human brain connectome. Annu Rev Clin Psychol. 2011;7:113–40.

54. Achard S, Bullmore E. Efficiency and Cost of Economical Brain Functional Networks. PLOS Comput Biol. 2007 Feb 2;3(2):e17.

55. Hagler DJ, Hatton SeanN, Cornejo MD, Makowski C, Fair DA, Dick AS, et al. Image processing and analysis methods for the Adolescent Brain Cognitive Development Study. NeuroImage. 2019 Nov 15;202:116091.

56. Mamah D, Barch DM, Repovš G. Resting state functional connectivity of five neural networks in bipolar disorder and schizophrenia. J Affect Disord. 2013 Sep;150(2):601–9.

57. Repovš G, Barch DM. Working Memory Related Brain Network Connectivity in Individuals with Schizophrenia and Their Siblings. Front Hum Neurosci [Internet]. 2012 [cited 2020 May 10];6. Available from: http://journal.frontiersin.org/article/10.3389/fnhum.2012.00137/abstract

58. Repovs G, Csernansky JG, Barch DM. Brain Network Connectivity in Individuals with Schizophrenia and Their Siblings. Biol Psychiatry. 2011 May;69(10):967–73.

59. Van Dijk KRA, Hedden T, Venkataraman A, Evans KC, Lazar SW, Buckner RL. Intrinsic Functional Connectivity As a Tool For Human Connectomics: Theory, Properties, and Optimization. J Neurophysiol. 2010 Jan;103(1):297–321.

60. Ji JL, Spronk M, Kulkarni K, Repovš G, Anticevic A, Cole MW. Mapping the human brain’s cortical-subcortical functional network organization. NeuroImage. 2019 Jan;185:35–57.

61. Glasser MF, Coalson TS, Robinson EC, Hacker CD, Harwell J, Yacoub E, et al. A multi-modal parcellation of human cerebral cortex. Nature. 2016 Aug;536(7615): 171–8.

62. Blondel VD, Guillaume J-L, Lambiotte R, Lefebvre E. Fast unfolding of communities in large networks. J Stat Mech Theory Exp. 2008 Oct;2008(10):P10008.

63. Floares AG, Ferisgan M, Onita D, Ciuparu A, Calin GA, Manolache FB. The Smallest Sample Size for the Desired Diagnosis Accuracy. Int J Oncol Cancer Ther. 2017;2:13–9.

64. Mukherjee S, Tamayo P, Rogers S, Rifkin R, Engle A, Campbell C, et al. Estimating Dataset Size Requirements for Classifying DNA Microarray Data. J Comput Biol. 2003 Apr;10(2): 119–42.

65. Duda RO, Hart PE, Stork DG. Pattern Classification. John Wiley & Sons; 2012. 679 p.

66. Friedman JH. Greedy Function Approximation: A Gradient Boosting Machine. Ann Stat. 2001;29(5):1189–232.

67. Janssen RJ, Mourão-Miranda J, Schnack HG. Making Individual Prognoses in Psychiatry Using Neuroimaging and Machine Learning. Biol Psychiatry Cogn Neurosci Neuroimaging. 2018 Sep 1;3(9):798–808.

68. Bzdok D, Meyer-Lindenberg A. Machine Learning for Precision Psychiatry: Opportunities and Challenges. Biol Psychiatry Cogn Neurosci Neuroimaging. 2018 Mar 1;3(3):223–30.

69. Dwyer DB, Falkai P, Koutsouleris N. Machine Learning Approaches for Clinical Psychology and Psychiatry. Annu Rev Clin Psychol. 2018;14(1):91–118.

70. Iniesta R, Stahl D, McGuffin P. Machine learning, statistical learning and the future of biological research in psychiatry. Psychol Med. 2016 Sep;46(12):2455–65.

71. Cearns M, Hahn T, Baune BT. Recommendations and future directions for supervised machine learning in psychiatry. Transl Psychiatry. 2019 Oct 22;9(1): 1–12.

72. Rutledge RB, Chekroud AM, Huys QJ. Machine learning and big data in psychiatry: toward clinical applications. Curr Opin Neurobiol. 2019 Apr 1;55:152–9.

73. Chandler C, Foltz PW, Elvevåg B. Using Machine Learning in Psychiatry: The Need to Establish a Framework That Nurtures Trustworthiness. Schizophr Bull. 2020 Jan 4;46(1): 11–4.

74. Lundberg SM, Nair B, Vavilala MS, Horibe M, Eisses MJ, Adams T, et al. Explainable machine-learning predictions for the prevention of hypoxaemia during surgery. Nat Biomed Eng. 2018 Oct;2(10):749–60.

75. Ritchey M, Libby LA, Ranganath C. Chapter 3 - Cortico-hippocampal systems involved in memory and cognition: the PMAT framework. In: O’Mara S, Tsanov M, editors. Progress in Brain Research [Internet]. Elsevier; 2015 [cited 2021 Aug 6]. p. 45–64. (The Connected Hippocampus; vol. 219). Available from: https://www.sciencedirect.com/science/article/pii/S0079612315000588

76. Doll BB, Shohamy D, Daw ND. Multiple memory systems as substrates for multiple decision systems. Neurobiol Learn Mem. 2015 Jan 1;117:4–13.

77. Palomero-Gallagher N, Vogt BA, Schleicher A, Mayberg HS, Zilles K. Receptor architecture of human cingulate cortex: evaluation of the four-region neurobiological model. Hum Brain Mapp. 2009 Aug;30(8):2336–55.

78. Manza P, Tomasi D, Volkow ND. Subcortical Local Functional Hyperconnectivity in Cannabis Dependence. Biol Psychiatry Cogn Neurosci Neuroimaging. 2018;3(3):285–93.

79. Wu Y-F, Yang B. Gray matter changes in chronic heavy cannabis users: a voxel-level study using multivariate pattern analysis approach. Neuroreport. 2020 Dec 9;31(17):1236–41.

80. Cheng H, Skosnik P, Pruce B, Brumbaugh M, Vollmer J, Fridberg D, et al. Resting state functional magnetic resonance imaging reveals distinct brain activity in heavy cannabis users – a multi-voxel pattern analysis. J Psychopharmacol Oxf Engl. 2014 Nov;28(11): 1030–40.

81. Kahle EM, Veliz P, McCabe SE, Boyd CJ. Functional and structural social support, substance use and sexual orientation from a nationally representative sample of US adults. Addict Abingdon Engl. 2020 Mar;115(3):546–58.

82. Miles DR, van den Bree MBM, Pickens RW. Sex differences in shared genetic and environmental influences between conduct disorder symptoms and marijuana use in adolescents. Am J Med Genet. 2002 Mar 8;114(2):159–68.

83. Guxens M, Nebot M, Ariza C. Age and sex differences in factors associated with the onset of cannabis use: a cohort study. Drug Alcohol Depend. 2007 May 11;88(2–3):234–43.

84. Rusby JC, Light JM, Crowley R, Westling E. Influence of parent-youth relationship, parental monitoring, and parent substance use on adolescent substance use onset. J Fam Psychol JFP J Div Fam Psychol Am Psychol Assoc Div 43. 2018 Apr;32(3):310–20.

85. Farhat T, Simons-Morton B, Luk JW. Psychosocial correlates of adolescent marijuana use: variations by status of marijuana use. Addict Behav. 2011 Apr;36(4):404–7.

