## Supplement for "Explainable Machine Learning Analysis Reveals Gender Differences in the Phenotypic and Neurobiological Markers of Cannabis Use Disorder"

*Supplementary Table 1. Complete list of included phenotypic variables.*

| | HCP Database Variable Name | Description |
|---|---|---|
| 0 | MMSE_Score | Mini Mental Status Exam Total Score |
| 1 | PSQI_Score | PSQI: Sleep Total Score |
| 2 | PSQI_Comp1 | PSQI: Sleep Component 1 Score |
| 3 | PSQI_Comp2 | PSQI: Sleep Component 2 Score |
| 4 | PSQI_Comp3 | PSQI: Sleep Component 3 Score |
| 5 | PSQI_Comp4 | PSQI: Sleep Component 4 Score |
| 6 | PSQI_Comp5 | PSQI: Sleep Component 5 Score |
| 7 | PSQI_Comp6 | PSQI: Sleep Component 6 Score |
| 8 | PSQI_Comp7 | PSQI: Sleep Component 7 Score |
| 9 | PSQI_Min2Asleep | PSQI: Minutes to fall asleep (past month) |
| 10 | PSQI_AmtSleep | PSQI: Hours of sleep per night (past month) |
| 11 | PSQI_Latency30Min | PSQI: Sleep Trouble - Can't get to sleep within 30 minutes |
| 12 | PSQI_WakeUp | PSQI: Sleep Trouble - Wake up in middle of night or early morning |
| 13 | PSQI_Bathroom | PSQI: Sleep Trouble - Get up to use bathroom |
| 14 | PSQI_Breathe | PSQI: Sleep Trouble - Can't breathe comfortably |
| 15 | PSQI_Snore | PSQI: Sleep Trouble - Cough or snore loudly |
| 16 | PSQI_TooCold | PSQI: Sleep Trouble - Feel too cold |
| 17 | PSQI_TooHot | PSQI: Sleep Trouble - Feel too hot |
| 18 | PSQI_BadDream | PSQI: Sleep Trouble - Had bad dreams |
| 19 | PSQI_Pain | PSQI: Sleep Trouble - Have pain |
| 20 | PSQI_Other | PSQI: Sleep Trouble - Other |
| 21 | PSQI_Quality | PSQI: Describe overall sleep quality |
| 22 | PSQI_SleepMeds | PSQI: How often taken sleep medicine |
| 23 | PSQI_DayStayAwake | PSQI: How often trouble staying awake |
| 24 | PSQI_DayEnthusiasm | PSQI: How often trouble keeping up enthusiasm |
| 25 | PSQI_BedPtnrRmate | PSQI: Have bed partner or roommate |
| 26 | PSQI_GetUpTime | PSQI: Time get up in morning (past month) |
| 27 | PSQI_BedTime | PSQI: Usual bed time (past month) |
| 28 | PicSeq_AgeAdj | NIH Toolbox: Picture Sequence Memory Test (Age-Adjusted) |

| | | |
|---|---|---|
| 29 | CardSort_AgeAdj | NIH Toolbox: Dimensional Change Card Sort (Age-Adjusted) |
| 30 | Flanker_AgeAdj | NIH Toolbox: Flanker Inhibitory Control and Attention Test (Age-Adjusted) |
| 31 | PMAT24_A_CR | Penn Progressive Matrices: Number of Correct Responses |
| 32 | PMAT24_A_SI | Penn Progressive Matrices: Total Skipped Items |
| 33 | PMAT24_A_RTCR | Penn Progressive Matrices: Median RT Correct Responses |
| 34 | ReadEng_AgeAdj | NIH Toolbox: Oral Reading Recognition Test (Age-Adjusted) |
| 35 | PicVocab_AgeAdj | NIH Toolbox: Picture Vocabulary Test (Age-Adjusted) |
| 36 | ProcSpeed_AgeAdj | NIH Toolbox: Processing Speed (Age-Adjusted) |
| 37 | DDisc_SV_1mo_200 | Delay Discounting: Subjective Value 1 month $200 |
| 38 | DDisc_SV_6mo_200 | Delay Discounting: Subjective Value 6 months $200 |
| 39 | DDisc_SV_1yr_200 | Delay Discounting: Subjective Value 1 year $200 |
| 40 | DDisc_SV_3yr_200 | Delay Discounting: Subjective Value 3 years $200 |
| 41 | DDisc_SV_5yr_200 | Delay Discounting: Subjective Value 5 years $200 |
| 42 | DDisc_SV_10yr_200 | Delay Discounting: Subjective Value 10 years $200 |
| 43 | DDisc_SV_1mo_40K | Delay Discounting: Subjective Value 1 month $40K |
| 44 | DDisc_SV_6mo_40K | Delay Discounting: Subjective Value 6 months $40K |
| 45 | DDisc_SV_1yr_40K | Delay Discounting: Subjective Value 1 year $40K |
| 46 | DDisc_SV_3yr_40K | Delay Discounting: Subjective Value 3 years $40K |
| 47 | DDisc_SV_5yr_40K | Delay Discounting: Subjective Value 5 years $40K |
| 48 | DDisc_SV_10yr_40K | Delay Discounting: Subjective Value 10 years $40K |
| 49 | DDisc_AUC_200 | Delay Discounting: Area Under the Curve $200 |
| 50 | DDisc_AUC_40K | Delay Discounting: Area Under the Curve $40K |
| 51 | VSPLOT_TC | Variable Short Penn Line Orientation: Total Number Correct |
| 52 | VSPLOT_CRTE | Variable Short Penn Line Orientation: Median RT/Expected Number of Clicks Correct |
| 53 | VSPLOT_OFF | Variable Short Penn Line Orientation: Total Positions Off for All Trials |
| 54 | SCPT_TP | Short Penn Continuous Performance Test: True Positives |
| 55 | SCPT_TN | Short Penn Continuous Performance Test: True Negatives |
| 56 | SCPT_FP | Short Penn Continuous Performance Test: False Positives |
| 57 | SCPT_FN | Short Penn Continuous Performance Test: False Negatives |
| 58 | SCPT_TPRT | Short Penn Continuous Performance Test: Median RT True Positives |
| 59 | SCPT_SEN | Short Penn Continuous Performance Test: Sensitivity |
| 60 | SCPT_SPEC | Short Penn Continuous Performance Test: Specificity |
| 61 | SCPT_LRNR | Short Penn Continuous Performance Test: Longest Run of Non-Responses |
| 62 | IWRD_TOT | Penn Word Memory Test: Total Number of Correct Responses |
| 63 | IWRD_RTC | Penn Word Memory Test: Median RT Correct Responses |
| 64 | ListSort_AgeAdj | NIH Toolbox: List Sorting Working Memory Test (Age-Adjusted) |
| 65 | CogFluidComp_AgeAdj | NIH Toolbox: Cognition Fluid Composite (Age-Adjusted) |
| 66 | CogEarlyComp_AgeAdj | NIH Toolbox: Cognition Early Childhood Composite (Age-Adjusted) |
| 67 | CogTotalComp_AgeAdj | NIH Toolbox: Cognition Total Composite Score (Age-Adjusted) |
| 68 | CogCrystalComp_AgeAdj | NIH Toolbox: Cognition Crystallized Composite (Age-Adjusted) |
| 69 | ER40_CR | Penn Emotion Recognition Test: Number of Correct Responses |

| | | |
|---|---|---|
| 70 | ER40_CRT | Penn Emotion Recognition Test: Correct Responses Median RT (ms) |
| 71 | ER40ANG | Penn Emotion Recognition Test: Number of Correct Anger |
| 72 | ER40FEAR | Penn Emotion Recognition Test: Number of Correct Fear |
| 73 | ER40HAP | Penn Emotion Recognition Test: Number of Correct Happy |
| 74 | ER40NOE | Penn Emotion Recognition Test: Number of Correct Neutral |
| 75 | ER40SAD | Penn Emotion Recognition Test: Number of Correct Sad |
| 76 | AngAffect_Unadj | NIH Toolbox: Anger-Affect Survey Score |
| 77 | AngHostil_Unadj | NIH Toolbox: Anger-Hostility Survey Score |
| 78 | AngAggr_Unadj | NIH Toolbox: Anger-Physical Aggression Survey Score |
| 79 | FearAffect_Unadj | NIH Toolbox: Fear-Affect Arousal Survey Score |
| 80 | FearSomat_Unadj | NIH Toolbox: Fear-Somatic Arousal Survey Score |
| 81 | Sadness_Unadj | NIH Toolbox: Sadness Survey Score |
| 82 | LifeSatisf_Unadj | NIH Toolbox: General Life Satisfaction Survey Score |
| 83 | MeanPurp_Unadj | NIH Toolbox: Meaning and Purpose Survey Score |
| 84 | PosAffect_Unadj | NIH Toolbox: Positive Affect Survey Score |
| 85 | Friendship_Unadj | NIH Toolbox: Friendship Survey Score |
| 86 | Loneliness_Unadj | NIH Toolbox: Loneliness Survey Score |
| 87 | PercHostil_Unadj | NIH Toolbox: Perceived Hostility Survey Score |
| 88 | PercReject_Unadj | NIH Toolbox: Perceived Rejection Survey Score |
| 89 | EmotSupp_Unadj | NIH Toolbox: Emotional Support Survey Score |
| 90 | InstruSupp_Unadj | NIH Toolbox: Instrumental Support Survey Score |
| 91 | PercStress_Unadj | NIH Toolbox: Perceived Stress Survey Score |
| 92 | SelfEff_Unadj | NIH Toolbox: Self-Efficacy Survey Score |
| 93 | Emotion_Task_Acc | fMRI OVERALL Emotion Task Accuracy |
| 94 | Emotion_Task_Median_RT | fMRI OVERALL Emotion Task RT |
| 95 | Emotion_Task_Face_Acc | fMRI Emotion Task FACE Accuracy |
| 96 | Emotion_Task_Face_Median_RT | fMRI Emotion Task FACE Median RT |
| 97 | Emotion_Task_Shape_Acc | fMRI Emotion Task SHAPE Accuracy |
| 98 | Emotion_Task_Shape_Median_RT | fMRI Emotion Task SHAPE Median RT |
| 99 | Gambling_Task_Perc_Larger | fMRI Gambling Task Overall Percentage 'Larger' |
| 100 | Gambling_Task_Perc_Smaller | fMRI Gambling Task Overall Percentage 'Smaller' |
| 101 | Gambling_Task_Perc_NLR | fMRI Gambling Task Overall Percentage No Logged Response |
| 102 | Gambling_Task_Median_RT_Larger | fMRI Gambling Task Overall RT 'Larger' |
| 103 | Gambling_Task_Median_RT_Smaller | fMRI Gambling Task Overall RT 'Smaller' |
| 104 | Gambling_Task_Reward_Perc_Larger | fMRI Gambling Task Percentage 'Larger' in Reward |
| 105 | Gambling_Task_Reward_Median_RT_Larger | fMRI Gambling Task Median RT 'Larger' in Reward |
| 106 | Gambling_Task_Reward_Perc_Smaller | fMRI Gambling Task Percentage 'Smaller' in Reward |
| 107 | Gambling_Task_Reward_Median_RT_Smaller | fMRI Gambling Task Median RT 'Smaller' in Reward |
| 108 | Gambling_Task_Reward_Perc_NLR | fMRI Gambling Task Percentage No Logged Response in Reward |
| 109 | Gambling_Task_Punish_Perc_Larger | fMRI Gambling Task Percentage 'Larger' in Punish |
| 110 | Gambling_Task_Punish_Median_RT_Larger | fMRI Gambling Task Median RT 'Larger' in Punish |

| | | |
|---|---|---|
| 111 | Gambling_Task_Punish_Perc_Smaller | fMRI Gambling Task Percentage 'Smaller' in Punish |
| 112 | Gambling_Task_Punish_Median_RT_Smaller | fMRI Gambling Task Median RT 'Smaller' in Punish |
| 113 | Gambling_Task_Punish_Perc_NLR | fMRI Gambling Task Percentage No Logged Response in Punish |
| 114 | Language_Task_Acc | fMRI Language Task OVERALL Accuracy |
| 115 | Language_Task_Median_RT | fMRI Language Task OVERALL Median RT |
| 116 | Language_Task_Story_Acc | fMRI Language Task STORY Accuracy |
| 117 | Language_Task_Story_Median_RT | fMRI Language Task STORY Median RT |
| 118 | Language_Task_Story_Avg_Difficulty_Level | fMRI Language Task STORY Difficulty Level |
| 119 | Language_Task_Math_Acc | fMRI Language Task MATH Accuracy |
| 120 | Language_Task_Math_Median_RT | fMRI Language Task MATH Median RT |
| 121 | Language_Task_Math_Avg_Difficulty_Level | fMRI Language Task MATH Difficulty Level |
| 122 | Relational_Task_Acc | fMRI Relational Task OVERALL Accuracy |
| 123 | Relational_Task_Median_RT | fMRI Relational Task OVERALL RT |
| 124 | Relational_Task_Match_Acc | fMRI Relational Task MATCH Accuracy |
| 125 | Relational_Task_Match_Median_RT | fMRI Relational Task MATCH Median RT |
| 126 | Relational_Task_Rel_Acc | fMRI Relational Task RELATIONAL block (REL) Accuracy |
| 127 | Relational_Task_Rel_Median_RT | fMRI Relational Task RELATIONAL block (REL) Median RT |
| 128 | Social_Task_Perc_Random | fMRI Social Task Overall Percentage 'Random' |
| 129 | Social_Task_Perc_TOM | fMRI Social Task Overall Percentage 'TOM' |
| 130 | Social_Task_Perc_Unsure | fMRI Social Task Overall Percentage 'Unsure' |
| 131 | Social_Task_Perc_NLR | fMRI Social Task Overall Percentage No Logged Response |
| 132 | Social_Task_Median_RT_Random | fMRI Social Task Overall RT 'Random' |
| 133 | Social_Task_Median_RT_TOM | fMRI Social Task Overall RT 'TOM' |
| 134 | Social_Task_Median_RT_Unsure | fMRI Social Task Overall RT 'Unsure' |
| 135 | Social_Task_Random_Perc_Random | fMRI Social Task Percentage 'Random' in Random condition |
| 136 | Social_Task_Random_Median_RT_Random | fMRI Social Task Median RT 'Random' in Random condition |
| 137 | Social_Task_Random_Perc_TOM | fMRI Social Task Percentage 'TOM' in Random condition |
| 138 | Social_Task_Random_Median_RT_TOM | fMRI Social Task Median RT 'TOM' in Random condition |
| 139 | Social_Task_Random_Perc_Unsure | fMRI Social Task Percentage 'Unsure' in Random Condition |
| 140 | Social_Task_Random_Median_RT_Unsure | fMRI Social Task Median RT 'Unsure' in Random condition |
| 141 | Social_Task_Random_Perc_NLR | fMRI Social Task Percentage No Logged Response in Random condition |
| 142 | Social_Task_TOM_Perc_Random | fMRI Social Task Percentage 'Random' in Social (TOM) condition |
| 143 | Social_Task_TOM_Median_RT_Random | fMRI Social Task Median RT 'Random' in Social (TOM) condition |
| 144 | Social_Task_TOM_Perc_TOM | fMRI Social Task Percentage 'TOM' in Social (TOM) condition |
| 145 | Social_Task_TOM_Median_RT_TOM | fMRI Social Task Median RT 'TOM' in Social (TOM) condition |
| 146 | Social_Task_TOM_Perc_Unsure | fMRI Social Task Percentage 'Unsure' in Social (TOM) condition |
| 147 | Social_Task_TOM_Median_RT_Unsure | fMRI Social Task Median RT 'Unsure' in Social (TOM) condition |
| 148 | Social_Task_TOM_Perc_NLR | fMRI Social Task Percentage No Logged Response in Social (TOM) condition |
| 149 | WM_Task_Acc | fMRI Working Memory Task Overall Accuracy |
| 150 | WM_Task_Median_RT | fMRI Working Memory Task Overall RT |

| | | |
|---|---|---|
| 151 | WM_Task_2bk_Acc | fMRI Working Memory Task Accuracy for 2-back |
| 152 | WM_Task_2bk_Median_RT | fMRI Working Memory Task Median RT for 2-back |
| 153 | WM_Task_0bk_Acc | fMRI Working Memory Task Accuracy for 0-back |
| 154 | WM_Task_0bk_Median_RT | fMRI Working Memory Task Median RT for 0-back |
| 155 | WM_Task_0bk_Body_Acc | fMRI Working Memory Task Accuracy for 0-back Body |
| 156 | WM_Task_0bk_Body_Acc_Target | fMRI Working Memory Task Accuracy for 0-back Body Targets |
| 157 | WM_Task_0bk_Body_Acc_Nontarget | fMRI Working Memory Task Accuracy for 0-back Body Nontargets |
| 158 | WM_Task_0bk_Face_Acc | fMRI Working Memory Task Accuracy for 0-back Face |
| 159 | WM_Task_0bk_Face_Acc_Target | fMRI Working Memory Task Accuracy for 0-back Face Targets |
| 160 | WM_Task_0bk_Face_ACC_Nontarget | fMRI Working Memory Task Accuracy for 0-back Face Nontargets |
| 161 | WM_Task_0bk_Place_Acc | fMRI Working Memory Task Accuracy for 0-back Place |
| 162 | WM_Task_0bk_Place_Acc_Target | fMRI Working Memory Task Accuracy for 0-back Place Targets |
| 163 | WM_Task_0bk_Place_Acc_Nontarget | fMRI Working Memory Task Accuracy for 0-back Place Nontargets |
| 164 | WM_Task_0bk_Tool_Acc | fMRI Working Memory Task Accuracy for 0-back Tool |
| 165 | WM_Task_0bk_Tool_Acc_Target | fMRI Working Memory Task Accuracy for 0-back Tool Targets |
| 166 | WM_Task_0bk_Tool_Acc_Nontarget | fMRI Working Memory Task Accuracy for 0-back Tool Nontargets |
| 167 | WM_Task_2bk_Body_Acc | fMRI Working Memory Task Accuracy for 2-back Body |
| 168 | WM_Task_2bk_Body_Acc_Target | fMRI Working Memory Task Accuracy for 2-back Body Targets |
| 169 | WM_Task_2bk_Body_Acc_Nontarget | fMRI Working Memory Task Accuracy for 2-back Body Nontargets |
| 170 | WM_Task_2bk_Face_Acc | fMRI Working Memory Task Accuracy for 2-back Face |
| 171 | WM_Task_2bk_Face_Acc_Target | fMRI Working Memory Task Accuracy for 2-back Face Targets |
| 172 | WM_Task_2bk_Face_Acc_Nontarget | fMRI Working Memory Task Accuracy for 2-back Face Nontargets |
| 173 | WM_Task_2bk_Place_Acc | fMRI Working Memory Task Accuracy for 2-back Place |
| 174 | WM_Task_2bk_Place_Acc_Target | fMRI Working Memory Task Accuracy for 2-back Place Targets |
| 175 | WM_Task_2bk_Place_Acc_Nontarget | fMRI Working Memory Task Accuracy for 2-back Place Nontargets |
| 176 | WM_Task_2bk_Tool_Acc | fMRI Working Memory Task Accuracy for 2-back Tool |
| 177 | WM_Task_2bk_Tool_Acc_Target | fMRI Working Memory Task Accuracy for 2-back Tool Targets |
| 178 | WM_Task_2bk_Tool_Acc_Nontarget | fMRI Working Memory Task Accuracy for 2-back Tool Nontargets |
| 179 | WM_Task_0bk_Body_Median_RT | fMRI Working Memory Task Median RT for 0-back Body |
| 180 | WM_Task_0bk_Body_Median_RT_Target | fMRI Working Memory Task Median RT for 0-back Body Targets |
| 181 | WM_Task_0bk_Body_Median_RT_Nontarget | fMRI Working Memory Task Median RT for 0-back Body Nontargets |
| 182 | WM_Task_0bk_Face_Median_RT | fMRI Working Memory Task Median RT for 0-back Face |
| 183 | WM_Task_0bk_Face_Median_RT_Target | fMRI Working Memory Task Median RT for 0-back Face Targets |
| 184 | WM_Task_0bk_Face_Median_RT_Nontarget | fMRI Working Memory Task Median RT for 0-back Face Nontargets |
| 185 | WM_Task_0bk_Place_Median_RT | fMRI Working Memory Task Median RT for 0-back Place |
| 186 | WM_Task_0bk_Place_Median_RT_Target | fMRI Working Memory Task Median RT for 0-back Place Targets |
| 187 | WM_Task_0bk_Place_Median_RT_Nontarget | fMRI Working Memory Task Median RT for 0-back Place Nontargets |
| 188 | WM_Task_0bk_Tool_Median_RT | fMRI Working Memory Task Median RT for 0-back Tool |
| 189 | WM_Task_0bk_Tool_Median_RT_Target | fMRI Working Memory Task Median RT for 0-back Tool Targets |
| 190 | WM_Task_0bk_Tool_Median_RT_Nontarget | fMRI Working Memory Task Median RT for 0-back Tool Nontargets |

| | | |
|---|---|---|
| 191 | WM_Task_2bk_Body_Median_RT | fMRI Working Memory Task Median RT for 2-back Body |
| 192 | WM_Task_2bk_Body_Median_RT_Target | fMRI Working Memory Task Median RT for 2-back Body Targets |
| 193 | WM_Task_2bk_Body_Median_RT_Nontarget | fMRI Working Memory Task Median RT for 2-back Body Nontargets |
| 194 | WM_Task_2bk_Face_Median_RT | fMRI Working Memory Task Median RT for 2-back Face |
| 195 | WM_Task_2bk_Face_Median_RT_Target | fMRI Working Memory Task Medain RT for 2-back Face Targets |
| 196 | WM_Task_2bk_Face_Median_RT_Nontarget | fMRI Working Memory Task Median RT for 2-back Face NonTargets |
| 197 | WM_Task_2bk_Place_Median_RT | fMRI Working Memory Task Median RT for 2-back Place |
| 198 | WM_Task_2bk_Place_Median_RT_Target | fMRI Working Memory Task Median RT for 2-back Place Targets |
| 199 | WM_Task_2bk_Place_Median_RT_Nontarget | fMRI Working Memory Task Median RT for 2-back Place Nontargets |
| 200 | WM_Task_2bk_Tool_Median_RT | fMRI Working Memory Task Median RT for 2-back Tool |
| 201 | WM_Task_2bk_Tool_Median_RT_Target | fMRI Working Memory Task Median RT for 2-back Tool Targets |
| 202 | WM_Task_2bk_Tool_Median_RT_Nontarget | fMRI Working Memory Task Median RT for 2-back Tool Nontargets |
| 203 | NEOFAC_A | NEO-FFI Agreeableness |
| 204 | NEOFAC_O | NEO-FFI Openness to Experience |
| 205 | NEOFAC_C | NEO-FFI Conscientiousness |
| 206 | NEOFAC_N | NEO-FFI Neuroticism |
| 207 | NEOFAC_E | NEO-FFI Extraversion |
| 208 | Age_in_Yrs | Age in Years |
| 209 | Gender_F | Female Gender |
| 210 | Race_Am. Indian/Alaskan Nat. | Race: Am. Indian/Alaskan Nat. |
| 211 | Race_Asian/Nat. Hawaiian/Othr Pacific Is. | Race: Asian/Nat. Hawaiian/Other Pacific Is. |
| 212 | Race_Black or African Am. | Race: Black/African American |
| 213 | Race_More than one | Race: More than one |
| 214 | Race_Unknown or Not Reported | Race: Unknown/Not Reported |
| 215 | Race_White | Race: White |
| 216 | Ethnicity_Hispanic/Latino | Ethnicity: Hispanic/Latino |
| 217 | Ethnicity_Not Hispanic/Latino | Ethnicity: Not Hispanic/Latino |
| 218 | Ethnicity_Unknown or Not Reported | Ethnicity: Unknown/Not Reported |
| 219 | ZygositySR_ | ZygositySR: Missing |
| 220 | ZygositySR_MZ | ZygositySR: MZ |
| 221 | ZygositySR_NotMZ | ZygositySR: Not MZ |
| 222 | ZygositySR_NotTwin | ZygositySR: Not Twin |
| 223 | ZygosityGT_ | ZygosityGT: Missing |
| 224 | ZygosityGT_DZ | ZygosityGT: DZ |
| 225 | ZygosityGT_MZ | ZygosityGT: MZ |
| 226 | TestRetestInterval | Test Retest Interval |
| 227 | Handedness | Handedness |
| 228 | SSAGA_Employ | SSAGA: Employment Status |
| 229 | SSAGA_Income | SSAGA: Household Income |
| 230 | SSAGA_Educ | SSAGA: Education |

| | | |
|---|---|---|
| 231 | SSAGA_InSchool | SSAGA: Still in School |
| 232 | SSAGA_Rlshp | SSAGA: Relationship Status |
| 233 | SSAGA_MOBorn | SSAGA: Missouri Born |
| 234 | FamHist_Moth_Scz | Mother Schizophrenia or Psychosis |
| 235 | FamHist_Fath_Scz | Father Schizophrenia or Psychosis |
| 236 | FamHist_Moth_Dep | Mother Depression |
| 237 | FamHist_Fath_Dep | Father Depression |
| 238 | FamHist_Moth_BP | Mother Bipolar Disorder |
| 239 | FamHist_Fath_BP | Father Bipolar Disorder |
| 240 | FamHist_Moth_Anx | Mother Anxiety Needing Treatment |
| 241 | FamHist_Fath_Anx | Father Anxiety Needing Treatment |
| 242 | FamHist_Moth_DrgAlc | Mother Drug or Alcohol Problems |
| 243 | FamHist_Fath_DrgAlc | Father Drug or Alcohol Problems |
| 244 | FamHist_Moth_Alz | Mother Alzheimer's or Dementia |
| 245 | FamHist_Fath_Alz | Father Alzheimer's or Dementia |
| 246 | FamHist_Moth_PD | Mother Parkinson's Disease |
| 247 | FamHist_Fath_PD | Father Parkinson's Disease |
| 248 | FamHist_Moth_TS | Mother Tourette's Syndrome |
| 249 | FamHist_Fath_TS | Father Tourette's Syndrome |
| 250 | FamHist_Moth_None | Mother None of the Above |
| 251 | FamHist_Fath_None | Father None of the Above |
| 252 | ASR_Anxd_Pct | ASR: Anxious/Depressed (Gender and Age-Adjusted Percentile) |
| 253 | ASR_Witd_T | ASR: Withdrawn (Gender and Age-Adjusted T-score) |
| 254 | ASR_Soma_T | ASR: Somatic Complaints (Gender and Age-Adjusted T-score) |
| 255 | ASR_Thot_T | ASR: Thought Problems (Gender and Age-Adjusted T-score) |
| 256 | ASR_Attn_T | ASR: Attention Problems (Gender and Age-Adjusted T-score) |
| 257 | ASR_Aggr_T | ASR: Aggressive Behavior (Gender and Age-Adjusted T-score) |
| 258 | ASR_Intr_T | ASR: Intrusive (Gender and Age-Adjusted T-score) |
| 259 | ASR_Intn_T | ASR: Internalizing (Gender and Age-Adjusted T-score) |
| 260 | ASR_Extn_T | ASR: Externalizing (Gender and Age-Adjusted T-score) |
| 261 | ASR_TAO_Sum | ASR: Sum of Thought, Attention, and Other Problems Raw Score |
| 262 | ASR_Totp_T | ASR: Total Problems (Gender and Age-Adjusted T-score) |
| 263 | DSM_Depr_T | ASR DSM: Depressive Problems (Gender and Age-Adjusted T-score) |
| 264 | DSM_Anxi_T | ASR DSM: Anxiety Problems (Gender and Age-Adjusted T-score) |
| 265 | DSM_Somp_T | ASR DSM: Somatic Problems (Gender and Age-Adjusted T-score) |
| 266 | DSM_Avoid_T | ASR DSM: Avoidant Personality Problems (Gender and Age-Adjusted T-score) |
| 267 | DSM_Adh_T | ASR DSM: AD/H Problems (Gender and Age-Adjusted T-score) |
| 268 | DSM_Antis_T | ASR DSM: Antisocial Personality Problems (Gender and Age-Adjusted T-score) |
| 269 | SSAGA_ChildhoodConduct | SSAGA: Childhood Conduct Problems |
| 270 | SSAGA_PanicDisorder | SSAGA: Panic Disorder |
| 271 | SSAGA_Agoraphobia | SSAGA: Agoraphobia |

| | | |
|---|---|---|
| 272 | SSAGA_Depressive_Ep | SSAGA: Major Depressive Episode |
| 273 | SSAGA_Depressive_Sx | SSAGA: Number of Depressive Symptoms |

*Supplementary Table 2. Best performing hyperparameters for each model.*

| | Learning rate | Max depth | Subsample |
|---|---|---|---|
| Phenotypic+FreeSurfer 1+ | 0.01 | 10 | 0.6 |
| Phenotypic+FreeSurfer 10+ | 0.20 | 8 | 0.8 |
| Phenotypic+FreeSurfer 100+ | 0.01 | 12 | 0.6 |
| Phenotypic+FreeSurfer 1000+ | 0.02 | 6 | 0.6 |
| Phenotypic+FreeSurfer Dependence | 0.05 | 8 | 1.0 |
| Phenotypic+Global Efficiency 1+ | 0.01 | 12 | 0.6 |
| Phenotypic+Global Efficiency 10+ | 0.01 | 8 | 1.0 |
| Phenotypic+Global Efficiency 100+ | 0.20 | 4 | 0.6 |
| Phenotypic+Global Efficiency 1000+ | 0.10 | 6 | 1.0 |
| Phenotypic+Global Efficiency Dependence | 0.10 | 8 | 0.8 |

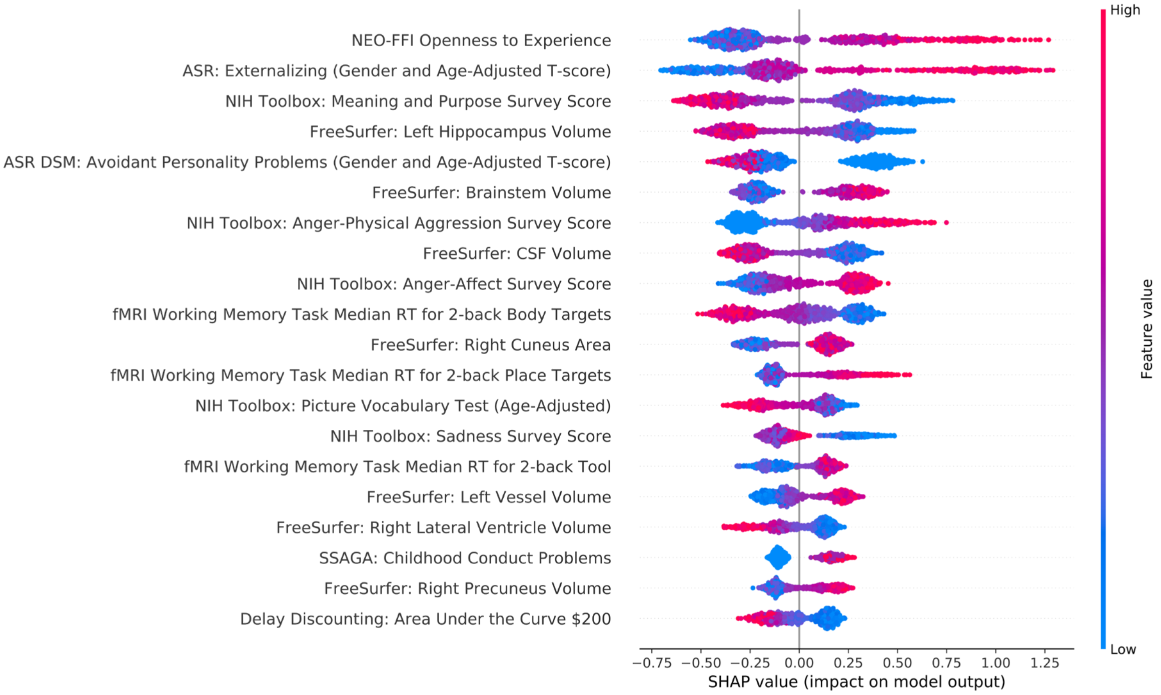

*Figure S1. SHAP factor ranking for the phenotypic+Freesurfer model classifying 1+ lifetime cannabis use. Factors are ranked in order of greatest average SHAP value, which indicates the importance of that factor. Individual points represent the model output for each individual in the sample. The position of a dot on the x-axis represents the impact of the observed factor on the model output for the individual. More positive SHAP values on the x-axis indicate that the observed factor pushed the classification closer towards cannabis dependence, whereas more negative SHAP values indicate that the factor pushed the classification away from cannabis dependence.The color of the individual dots represent the value of the observed measurements, with blue indicating lower and red higher values.*

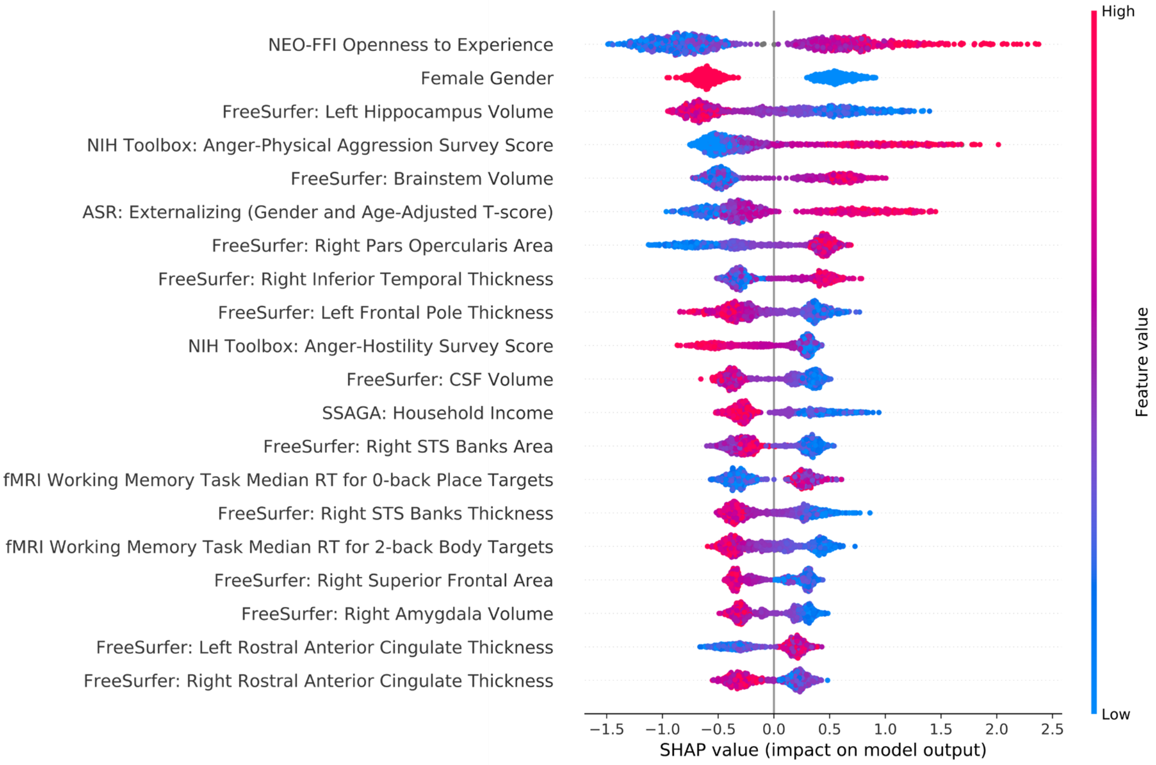


*Figure S1. SHAP factor ranking for the phenotypic+Freesurfer model classifying 10+ lifetime cannabis uses. Factors are ranked in order of greatest average SHAP value, which indicates the importance of the factor. Individual points represent the model output for each individual in the sample. The position of a dot on the x-axis represents the impact of the observed factor on the model output for the individual. More positive SHAP values on the x-axis indicate that the observed factor pushed the classification closer towards cannabis dependence, whereas more negative SHAP values indicate that the factor pushed the classification away from cannabis dependence.The color of the individual dots represent the value of the observed measurements, with blue indicating lower and red higher values.*

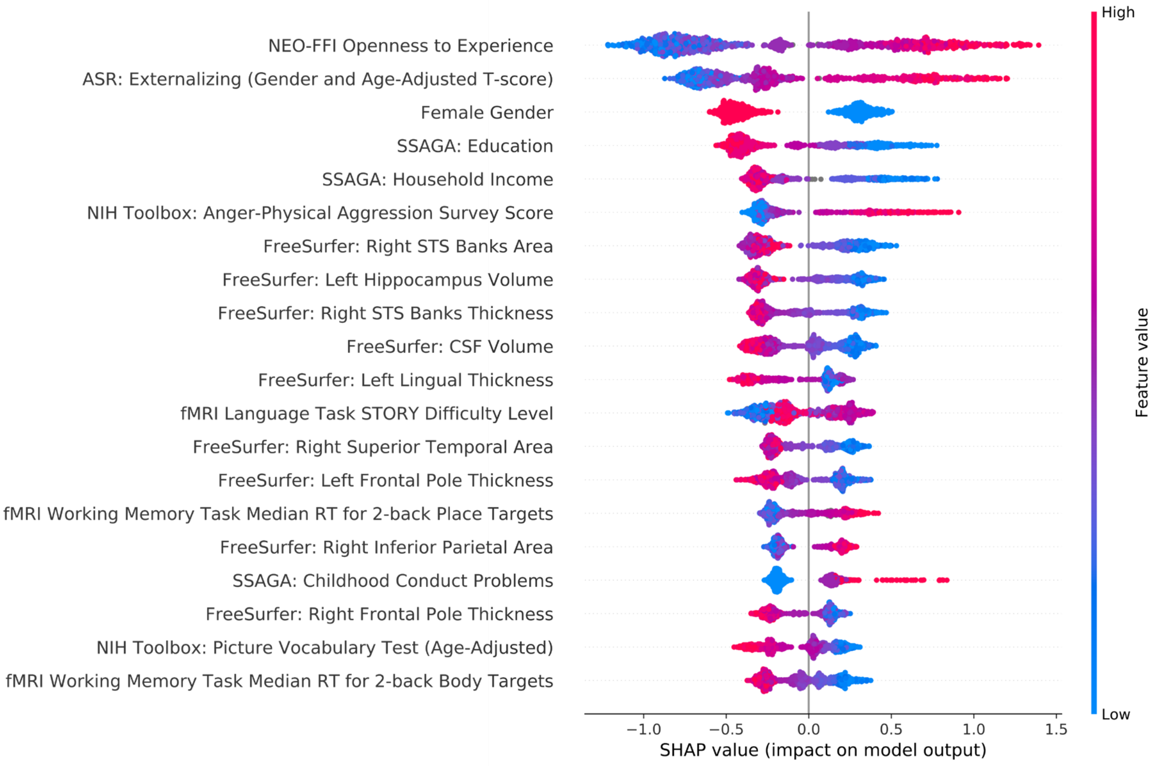


*Figure S1. SHAP factor ranking for the phenotypic+Freesurfer model classifying 100+ lifetime cannabis uses. Factors are ranked in order of greatest average SHAP value, which indicates the importance of the factor. Individual points represent the model output for each individual in the sample. The position of a dot on the x-axis represents the impact of the observed factor on the model output for the individual. More positive SHAP values on the x-axis indicate that the observed factor pushed the classification closer towards cannabis dependence, whereas more negative SHAP values indicate that the factor pushed the classification away from cannabis dependence.The color of the individual dots represent the value of the observed measurements, with blue indicating lower and red higher values.*

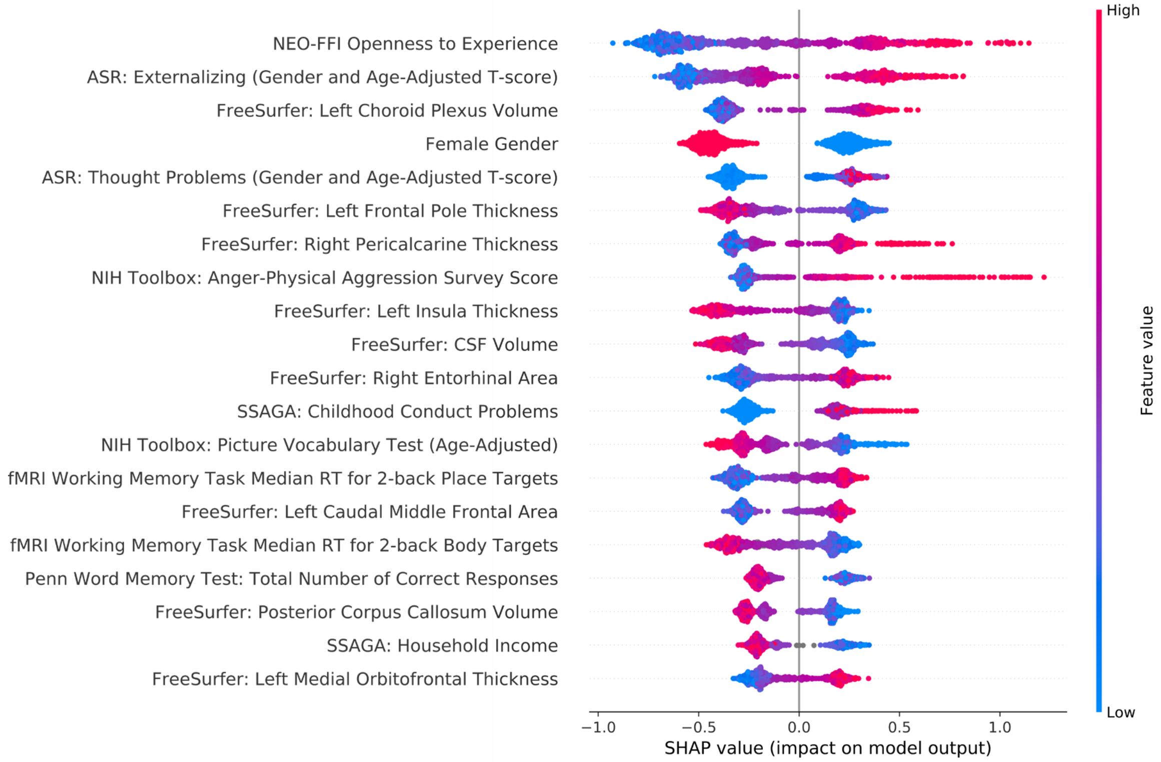


*Figure S1. SHAP factor ranking for the phenotypic+Freesurfer model classifying 1000+ lifetime cannabis uses. Factors are ranked in order of greatest average SHAP value, which indicates the importance of the factor. Individual points represent the model output for each individual in the sample. The position of a dot on the x-axis represents the impact of the observed factor on the model output for the individual. More positive SHAP values on the x-axis indicate that the observed factor pushed the classification closer towards cannabis dependence, whereas more negative SHAP values indicate that the factor pushed the classification away from cannabis dependence.The color of the individual dots represent the value of the observed measurements, with blue indicating lower and red higher values.*

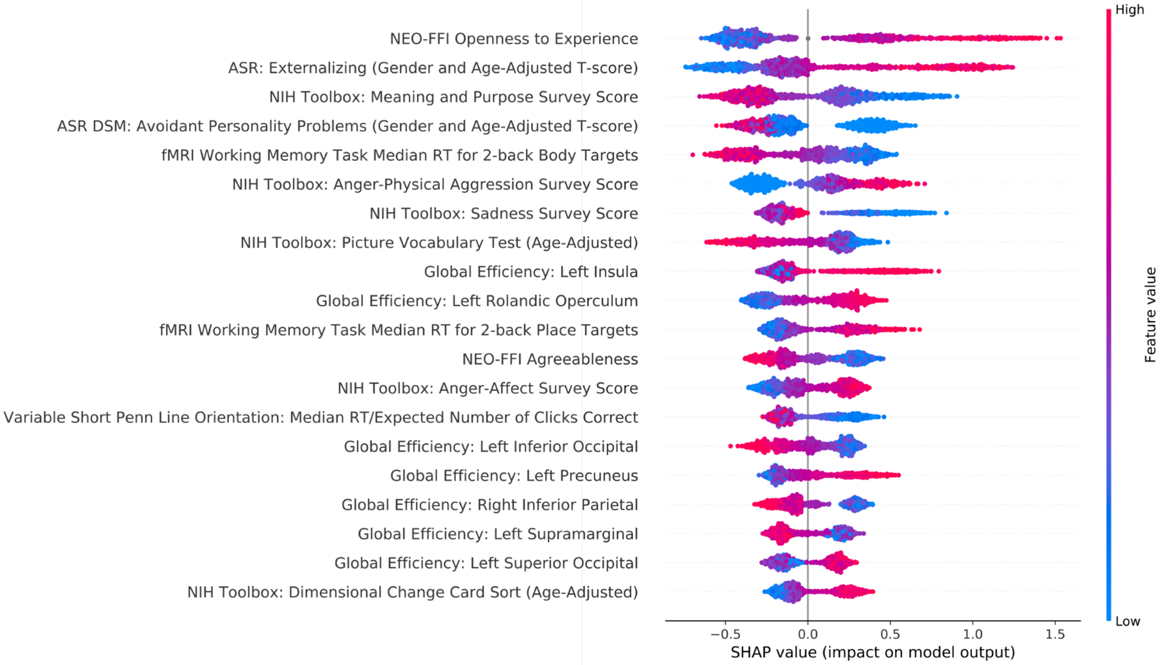


*Figure S1. SHAP factor ranking for the phenotypic+Global Efficiency model classifying 1+ lifetime cannabis uses. Factors are ranked in order of greatest average SHAP value, which indicates the importance of the factor. Individual points represent the model output for each individual in the sample. The position of a dot on the x-axis represents the impact of the observed factor on the model output for the individual. More positive SHAP values on the x-axis indicate that the observed factor pushed the classification closer towards cannabis dependence, whereas more negative SHAP values indicate that the factor pushed the classification away from cannabis dependence.The color of the individual dots represent the value of the observed measurements, with blue indicating lower and red higher values.*

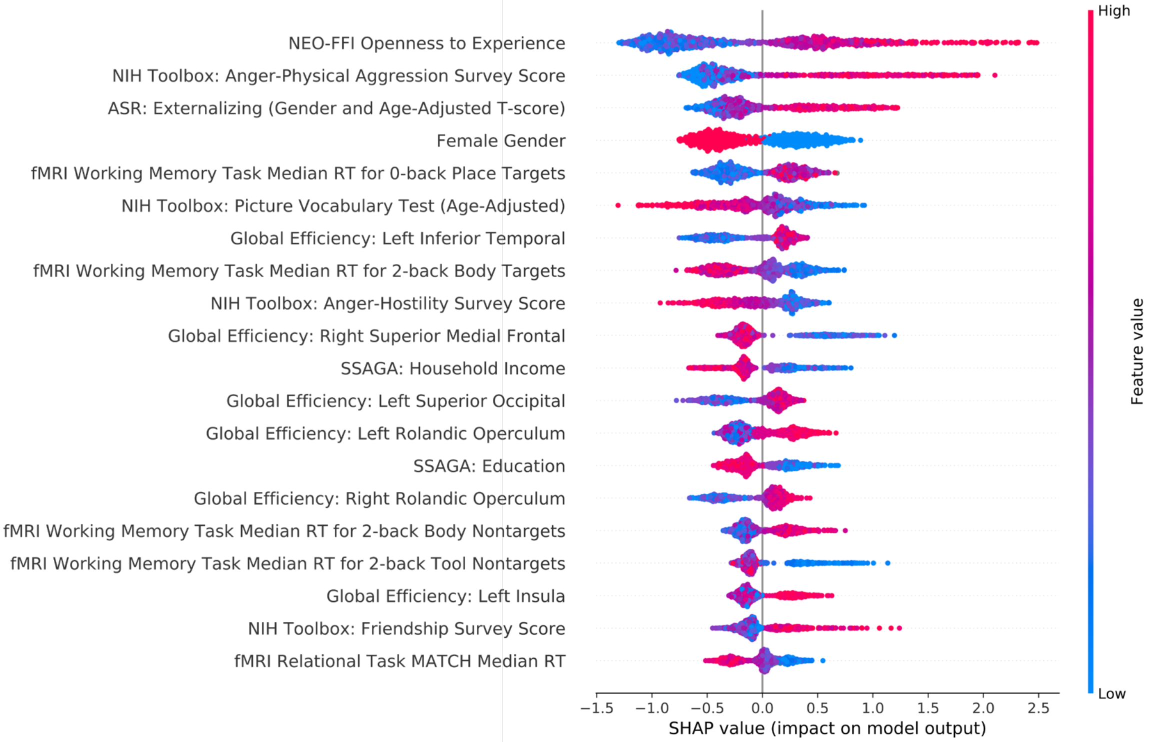


*Figure S1. SHAP factor ranking for the phenotypic+Global Efficiency model classifying 10+ lifetime cannabis uses. Factors are ranked in order of greatest average SHAP value, which indicates the importance of the factor. Individual points represent the model output for each individual in the sample. The position of a dot on the x-axis represents the impact of the observed factor on the model output for the individual. More positive SHAP values on the x-axis indicate that the observed factor pushed the classification closer towards cannabis dependence, whereas more negative SHAP values indicate that the factor pushed the classification away from cannabis dependence.The color of the individual dots represent the value of the observed measurements, with blue indicating lower and red higher values.*

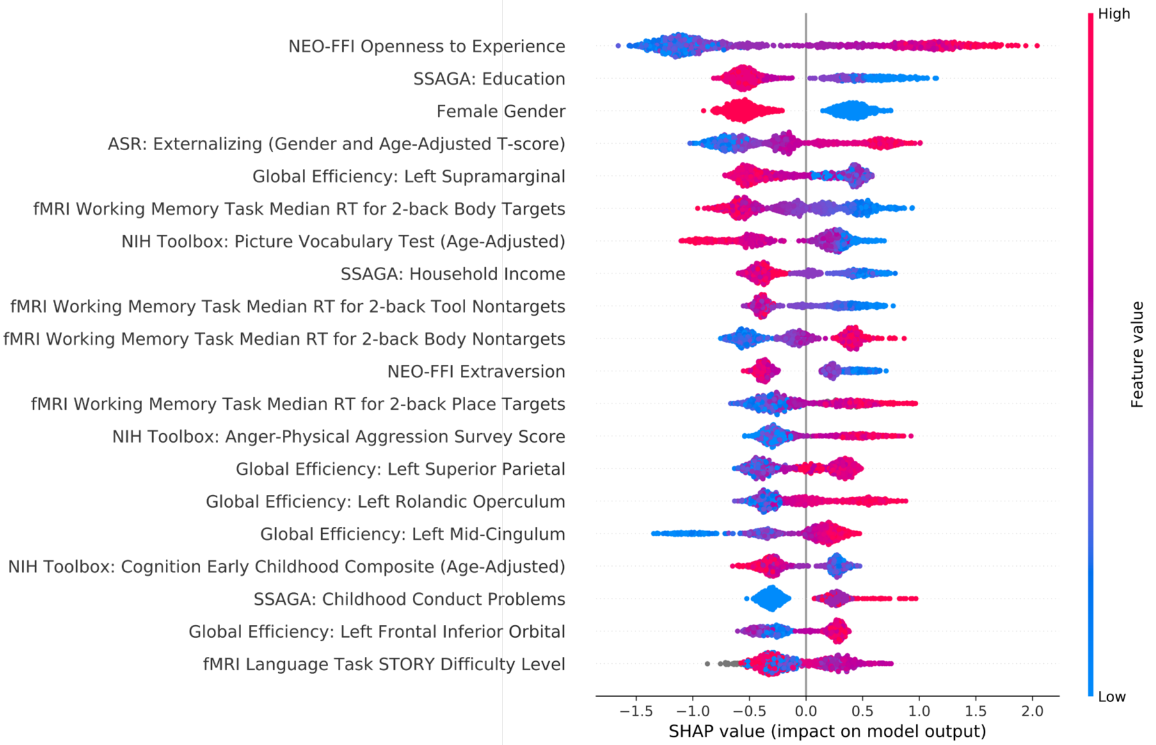


*Figure S1. SHAP factor ranking for the phenotypic+Global Efficiency model classifying 100+ lifetime cannabis uses. Factors are ranked in order of greatest average SHAP value, which indicates the importance of the factor. Individual points represent the model output for each individual in the sample. The position of a dot on the x-axis represents the impact of the observed factor on the model output for the individual. More positive SHAP values on the x-axis indicate that the observed factor pushed the classification closer towards cannabis dependence, whereas more negative SHAP values indicate that the factor pushed the classification away from cannabis dependence.The color of the individual dots represent the value of the observed measurements, with blue indicating lower and red higher values.*

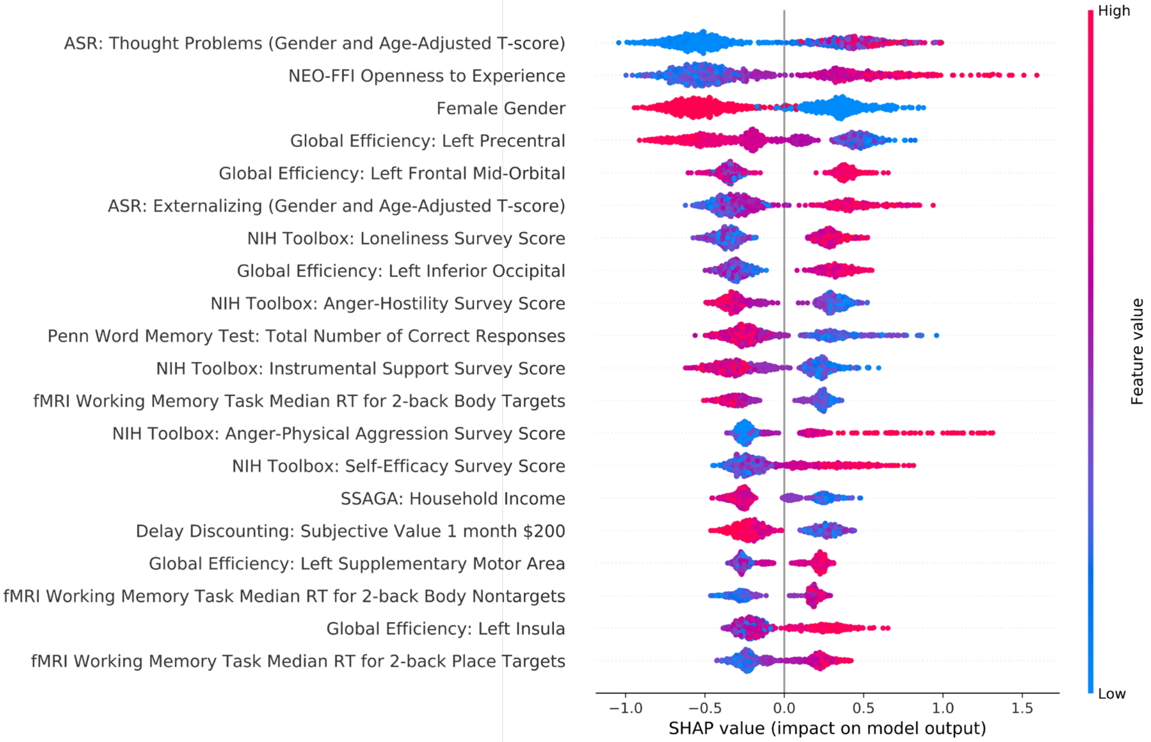


*Figure S1. SHAP factor ranking for the phenotypic+Global Efficiency model classifying 1000+ lifetime cannabis uses. Factors are ranked in order of greatest average SHAP value, which indicates the importance of the factor. Individual points represent the model output for each individual in the sample. The position of a dot on the x-axis represents the impact of the observed factor on the model output for the individual. More positive SHAP values on the x-axis indicate that the observed factor pushed the classification closer towards cannabis dependence, whereas more negative SHAP values indicate that the factor pushed the classification away from cannabis dependence.The color of the individual dots represent the value of the observed measurements, with blue indicating lower and red higher values.*
